# Gut bacterial lactate stimulates lung epithelial mitochondria and exacerbates acute lung injury

**DOI:** 10.1101/2025.03.24.645052

**Authors:** Vaibhav Upadhyay, Edwin F. Ortega, Luis A. Ramirez Hernandez, Margaret Alexander, Gagandeep Kaur, Kai Trepka, Rachel R. Rock, Rafaella T. Shima, W. Colby Cheshire, Narges Alipanah-Lechner, Carolyn S. Calfee, Michael A. Matthay, Joyce V. Lee, Andrei Goga, Isha H. Jain, Peter J. Turnbaugh

## Abstract

Acute respiratory distress syndrome (ARDS) is an often fatal critical illness where lung epithelial injury leads to intrapulmonary fluid accumulation. ARDS became widespread during the COVID-19 pandemic, motivating a renewed effort to understand the complex etiology of this disease. Rigorous prior work has implicated lung endothelial and epithelial injury in response to an insult such as bacterial infection; however, the impact of microorganisms found in other organs on ARDS remains unclear. Here, we use a combination of gnotobiotic mice, cell culture experiments, and re-analyses of a large metabolomics dataset from ARDS patients to reveal that gut bacteria impact lung cellular respiration by releasing metabolites that alter mitochondrial activity in lung epithelium. Colonization of germ-free mice with a complex gut microbiota stimulated lung mitochondrial gene expression. A single human gut bacterial species, *Bifidobacterium adolescentis,* was sufficient to replicate this effect, leading to a significant increase in mitochondrial membrane potential in lung epithelial cells. We then used genome sequencing and mass spectrometry to confirm that *B. adolescentis* produces *L*-lactate, which was sufficient to increase mitochondrial activity in lung epithelial cells. Finally, we found that serum lactate was significantly associated with disease severity in patients with ARDS from the Early Assessment of Renal and Lung Injury (EARLI) cohort. Together, these results emphasize the importance of more broadly characterizing the microbial etiology of ARDS and other lung diseases given the ability of gut bacterial metabolites to remotely control lung cellular respiration. Our discovery of a single bacteria-metabolite pair provides a *proof-of-concept* for systematically testing other microbial metabolites and a mechanistic biomarker that could be pursued in future clinical studies. Furthermore, our work adds to the growing literature linking the microbiome to mitochondrial function, raising intriguing questions as to the bidirectional communication between our endo- and ecto-symbionts.

## INTRODUCTION

Acute respiratory distress syndrome (ARDS) is a fatal and prevalent disease in critically ill patients^1^. ARDS commonly occurs due to viral or bacterial pneumonia, though is also caused by nonpulmonary sepsis, aspiration of gastric contents, and major trauma^2^. ARDS is characterized by the accumulation of protein-rich fluid in the alveolar air space that is cleared by epithelial cells^3,4^. Epithelial cells comprise most of the lung’s surface area^5^, and epithelial metabolism, including mitochondrial activity, is impaired in ARDS^6–8^.

ARDS was the number one cause of mortality for intensive care unit (ICU) patients during the initial phases of the COVID-19 pandemic^9^ because SARS-CoV-2 was a novel driver of acute epithelial injury and lung inflammation^10^. While vaccination reduced the burden of ARDS caused by SARS-CoV-2, ARDS remains prevalent in 10% of critically ill patients, including 23% of mechanically ventilated patients^2,11,12^. The treatment of ARDS is based on lung protective ventilation^13^, and the lack of pharmacotherapies to treat ARDS contributed to why COVID-19 caused such a significant burden on the healthcare system. Work on ARDS will be essential to address the still ongoing exposures to respiratory viruses and prepare for future pandemics^14^.

In addition to viral infections, bacterial pulmonary infections are a common cause of ARDS modeled experimentally, from which much information has been gleaned^15^. Injured lung epithelium exhibits denudation from the basement membrane and cell death patterns consistent with both necrosis and apoptosis^16–19^. Mitochondrial dysfunction is common in critically ill patients^20^, and serum mitochondrial nucleic acid is correlated with the onset of ARDS^21^. Bacterial products are among the stimuli that cause epithelial injury and local inflammation. Intrapulmonary lipopolysaccharide (LPS) results in interalveolar fluid accumulation^22^. While LPS is directly injurious to epithelial cells, LPS administration also results in the local release of IL-17A and an expansion of T helper 17 (Th17) cells^23^. In addition to bacterial products, other direct causes of lung epithelial injury include acid, excess oxygen, and mechanical forces^1,2^.

While the lung is constantly exposed to microorganisms in the air, the contribution of the resident microbiome to ARDS and other lung diseases is not well defined. This may be because the lower airway has incredibly low microbial biomass that is difficult to distinguish from background contaminants^24^, and the degree to which lung tissue harbors a resident microbiota in the absence of overt disease remains controversial^25,26^. While bacterial isolates have been sequenced from the lung in settings like ARDS^27,28^, this is after the onset of disease which results in significant changes to the lung microenvironment. In contrast, the gastrointestinal tract harbors hundreds of trillions of microorganisms (the gut microbiota) whose metabolic products are continuously absorbed into general circulation and transported to lung tissue^29,30^.

Emerging data has hinted at a potential pathogenic role for the gut microbiome in lung diseases like ARDS. The gut microbiota exacerbates interalveolar fluid accumulation in oxygen-induced lung injury, a mouse model of ARDS^31–33^. Furthermore, the gut microbiota increases pulmonary edema in mice exposed to gastrointestinal ischemia-reperfusion injury, where both local and remote inflammation is diminished in germ-free (GF) mice^34^. The interpretation of these observations is complex because the microbiome plays a vital role in immune development^35^, which may also contribute to lung health. Furthermore, the gut microbiota protects gastrointestinal epithelial cells from injurious stimuli^36^, though whether gut bacteria impact epithelial cells in other tissue beds is an open area of investigation.

We sought to use paired studies in mice and human subjects to evaluate the impact of specific gut bacteria on lung pathophysiology. Colonization of GF mice with a prevalent and abundant gut Actinomycetota, *Bifidobacterium adolescentis*, stimulates mitochondrial activity in lung epithelium. Using a series of *in vivo* gnotobiotic experiments, complete genome sequencing, and metabolomic profiling we show *B. adolescentis* increases lung epithelial mitochondrial activity by producing the metabolite *L*-lactate. Finally, we provide support for translational relevance by demonstrating an association between lactate and disease severity in critically ill patients with ARDS from the Early Assessment of Renal and Lung Injury (EARLI) cohort^20,37^.

## METHODS

### Mice

All mouse experiments were approved by the UCSF Institutional Animal Care and Use Committee (IACUC approval number AN200526-00E). C57BL/6J male and female mice aged 6-10 weeks were obtained from the UCSF Gnotobiotics Core (gnotobiotics.ucsf.edu). Prior to colonization, mice were monitored weekly via PCR and culture to verify GF status. At the time of colonization, mice were gavaged with 200 μL *B. adolescentis* BD1 grown to stationary phase for 24 hours, 200 μL *E. lenta* DSM2243 was grown to stationary phase for 72 hours, or the cecal contents of a healthy SPF mouse resuspended in Brain Heart Infusion (BHI) media. After colonization, mice were housed in a gnotobiotic isolator (Class Biologically Clean) or Tecniplast isocages. Mice were colonized for 10-14 days. Colonization was confirmed via culture at the experimental endpoint.

### Bacterial Culture

Bacterial strains were cultured at 37°C in an anaerobic chamber (Coy Laboratory Products) (5% H_2_, 20% CO_2_, balanced with N_2_). *B*. *adolescentis* strain BD1 was isolated previously from an obese subject on a control diet^38^ and cultured in BHI supplemented with L-Cysteine-HCl (0.05% w/v) and resazurin (0.0001% w/v). *E. lenta* strain 2243 was cultured in BHI supplemented with L-Cysteine-HCl (0.05% w/v), hemin (5 μg/ml), L-Arginine (1%), vitamin K (1 μg/ml), and resazurin (0.0001% w/v) (BHI^CHAVR^).

### Isolation of single cell suspension from mouse lung

Mice were sacrificed and lungs were isolated and placed in 5 mL of Hank’s Balanced Salt Solution (HBSS) with 0.2 mg/mL Liberase TM and 25 μg/mL DNAse (Sigma, St. Louis, MO). Lung tissue was digested using the GentleMACS (Miltenyi Biotec) system using the pre-programmed “lung1” program. Samples were then incubated for 30 minutes at 37°C with gentle agitation on an incubating shaker (160 rotations per minute). After incubation, samples were processed on the GentleMacs using the pre-programmed “lung2” program and passed through 70 μM filters. Cell pellets were subjected to red blood cell lysis (RBC Lysis Buffer, Biolegend, San Diego, CA).

### Bulk RNA sequencing (RNA-seq) of host tissues

All samples were submitted to the UCSF Genomics Colab (colabs.ucsf.edu/genomics) for sequencing. Sequencing was conducted in three separate batches. The methods for each batch will be described here in order of their presentation in the manuscript.

In the first batch of sequencing, *B. adolescentis* and GF colonized mice were sacrificed with single-cell suspensions isolated from the entire lung and 10^6^ cells frozen at -80°C in 600 μL buffer RLT. Samples were then thawed and cells were lysed using pipetting. After cells were lysed, the Mini RNA Kit (Germantown, MD) was used to extract samples. One volume of 70% ethanol was then added and the resulting mixture loaded onto an RNeasy spin column. The column was then washed with buffers RW1 and RPE with two on-column DNAse treatments prior to sample elution. RNA quality was assessed using the 4200 Agilent TapeStation system (Santa Clara, CA). RNA libraries were prepared using a Universal Plus Ribosomal Depletion Kit (Tecan Männedorf, Switzerland). 30-100 ng of total RNA of each sample was subjected to RNA Fragmentation without Poly(A) selection. Next rRNA was removed using the FastSelect -rRNA/Globin Kit. A cDNA library was then created using 15 cycles of PCR amplification, and libraries were pooled at equimolar concentrations. Quantified library pools were diluted to 1 nM and sequenced on a MiniSeq to check for the quality of reads. Individual libraries were then normalized according to MiniSeq output reads by the percentage of coding genes and were sequenced on a HiSeq4000 single-ended 50-base pair libraries.

In the second batch of sequencing, GF mice, *B. adolescentis* colonized mice, *E. lenta* colonized mice, or mice conventionalized with the cecal contents of a healthy SPF mouse were sacrificed. Lung tissue was immediately harvested and frozen at -80°C. A total of 30 mg of frozen lung tissue was homogenized using a Rotastator homogenizer in 600 μL buffer RLT. After cells were homogenized, the Mini RNA Kit (Germantown, MD) was used to extract samples. One volume of 70% ethanol was then added and the resulting mixture loaded onto an RNeasy spin column. The column was then washed with buffers RW1 and RPE with two on-column DNAse treatments prior to sample elution. RNA quality was assessed using the 4200 Agilent TapeStation system (Santa Clara, CA). RNA libraries were prepared using a Universal Plus Ribosomal Depletion Kit (Tecan Männedorf, Switzerland). A total of 30-100 ng of total RNA of each sample was subjected to RNA Fragmentation without Poly(A) selection. Next rRNA was removed using the FastSelect -rRNA/Globin Kit. A cDNA library was then created using 15 cycles of PCR amplification, and libraries were pooled at equimolar concentrations. Quantified library pools were diluted to 1 nM and sequenced on a MiniSeq to check for the quality of reads. Individual libraries were then normalized according to MiniSeq output reads by the percentage of coding genes and were sequenced on a NovaSeqX using paired-end 100-based pair libraries. One GF mouse was an outlier on PCA and removed from downstream analysis.

In a third batch of sequencing, *B. adolescentis* colonized mice and GF mice were sacrificed with colon tissue isolated and colonic contents removed via scraping. Colon samples were frozen at -80°C in RNALater. Samples were thawed and 30 mg of the colon was resuspended in 600 μL buffer RLT. Colon tissue was then homogenized using a Rotastator homogenizer. One volume of 70% ethanol was then added and the resulting mixture was loaded onto an RNeasy spin column. The column was then washed with buffers RW1 and RPE with two on-column DNAse treatments prior to sample elution. RNA quality was assessed using the 4200 Agilent TapeStation system (Santa Clara, CA). RNA libraries were prepared using a Universal Plus Ribosomal Depletion Kit (Tecan Männedorf, Switzerland). 30-100 ng of total RNA of each sample was subjected to RNA Fragmentation without Poly(A) selection. Next rRNA was removed using the FastSelect -rRNA/Globin Kit. A cDNA library was then created using 15 cycles of PCR amplification, and libraries were pooled at equimolar concentrations. Quantified library pools were diluted to 1 nM and sequenced on a MiniSeq to check for the quality of reads. Individual libraries were then normalized according to MiniSeq output reads by the percentage of coding genes and were sequenced on a HiSeq4000 single-ended 50-base pair libraries.

### RNA-seq analysis

All output bulk-RNA sequencing reads were aligned to the mouse reference genome (GRCm38) and reads per gene matrix were counted using the Ensemble annotation build version 96 using STAR v2.7.5c^39^. Read counts were used as an input to DESeq2^40^ (v1.46.0) to test for differential gene expression between conditions using a Wald test. Genes passing a multiple test correction *p*-value of 0.1 (Benjamini-Hochberg method) were considered significant. The stats (version 3.6.2) package was used to generate Euclidean distance matrices and conduct principal coordinate plots using the prcomp and dist commands. Differentially expressed genes were annotated as having mitochondrial activity or contributing to one of the five mitochondrial electron transport complexes using the MitoCarta 3.0 database. To assess *Bifidobacterium* transcript in colon or lung tissue, all RNA reads from colon and lung tissue for *B. adolescentis* colonized and GF mice were aligned using Kraken 2. The resulting reads were filtered for the *Bifidobacterium* genera and displayed as a proportion of all input reads with comparisons made between the GF and *B. adolescentis* colonized group. Each bulk-RNA sequencing batch was analyzed independently.

### Single-cell RNA sequencing (scRNA-seq) from host tissues

Cell suspensions were stained with the Zombie Green™ Fixable Viability Kit (Sony Biotechnology San Jose, CA) and APC-conjugated anti-mouse CD45 (Clone QA17A26, Sony Biotechnology San Jose, CA). Cells were sorted on a Sony SH800S Fluorescence Activated Cell Sorter (Sony Biotechnology San Jose, CA) to separate live hematopoietic and live non-hematopoietic cells. 10,000 live CD45+ and live CD45-events were collected per mouse. Single-cell library preparation was then conducted immediately using a Chromium Next GEM Single Cell 3’ Kit v3.1 (10x Genomics Pleasanton, CA). Reagents were thawed and equilibrated to room temperature prior to loading the chip and preparation of a single-cell master mix. 70 μL of reverse transcription buffer and enzymes were combined with individual samples and loaded into the chip. Single Cell 3’ v.3.1 beads and partitioning oil were loaded into their respective ports. The 10x Gasket was then attached and the assembled chip was run on the Chromium Controller immediately. Samples were then subjected to cDNA amplification using the cDNA amplification reaction mix. Indexed libraries were sequenced using a NovaSeq 6000 (Illumina, Foster City, CA) instrument. The resulting reads were filtered and processed using cellranger 7.0.1 (10x Genomics Pleasanton, CA).

### scRNA-seq analysis

Output libraries were analyzed using Seurat (v5.1.0)^41^. Output 10x files were imported into R using the Read10X command. Files were then made into seurat objects. CD45+ and CD45-cell types were merged and anchored in a combined seurat object. Data was then scaled with UMAP run using a “pca” reduction and 1:30 dims. Cluster markers were exported and uploaded into the Mouse Cell Atlas in the Run-scMCA tab with the resulting best fit annotations applied to each cluster ^42^. Differentially expressed genes were measured between cluster based on colonization state, mitochondrial genes were selected using the MitoCarta 3.0 database. A module score of all mitochondrial genes differentially expressed by *B. adolescentis* in all clusters for either CD45+ or CD45-cells was calculated using the AddModuleScore command.

### Stool DNA extraction

Stool samples underwent DNA extraction conducted by the Microbial Genomics Core at the University of California, San Francisco. The DNA extraction process utilized a modified cetyltrimethylammonium bromide (CTAB) buffer-based protocol, as detailed in a published article^43^. Briefly, each frozen mice stool pellet sample was added into Lysing Matrix E tube (MP Biomedicals) containing 500 µL of 5% CTAB extraction buffer and subsequent incubation at 65 °C for 15 minutes. Following this, 500 µL phenol:chloroform:isoamyl alcohol (25:24:1) was added, followed by bead-beating at 5.5 m/s for 30 seconds and centrifugation at 16,000g for 5 minutes at 4 °C. The resulting aqueous phase (∼400 µL) was transferred to a new 2mL Eppendorf tube. Additional 5% CTAB extraction buffer (400 µL) was added to the fecal aliquot, repeating the steps to yield approximately 800 µL from repeated extractions. Chloroform was added in equal volume, mixed, and centrifuged (16,000g for 5 minutes). The resulting aqueous phase (∼500 µL) was transferred to another 2mL Eppendorf tube, combined with a 2-volume of 30% polyethylene glycol (PEG)/NaCl solution, and stored at 4 °C overnight to precipitate DNA. Subsequently, samples were centrifuged (3,000g for 60 minutes), washed twice with ice-cold 70% ethanol, and resuspended in 100 µL sterile water. A negative extraction control was included in sample processing to account for potential contaminants in reagents. The extracted DNA from each sample was quantified using the Qubit 2.0 Fluorometer with the dsDNA BR Assay Kit (Life Technologies, Grand Island, NY) and normalized to 5 ng/µL using QIAgility Liquid Handler.

### Amplicon library preparation and sequencing

The hypervariable region 4 (v4) of the 16S rRNA gene was amplified via PCR using 515F/806R primers. The reverse primers (806R) contained a unique barcode sequence to enable demultiplexing of pooled samples, and an adapter sequence that enables the amplicons to bind to the flow cell. Each sample was amplified in a 25 μl reaction using 0.025 U Takara Hot Start ExTaq (Takara Mirus Bio Inc., Madison, WI), 1X Takara buffer, 0.4 pmol/μl of F515 and R806 primers, 0.56 mg/ml of bovine serum albumin (BSA; Roche Applied Science, Indianapolis, IN), 200 μM of dNTPs, and 10 ng of template DNA. The PCR conditions consisted of an initial denaturation at 98 °C for 2 minutes, followed by 30 cycles at 98 °C for 20 seconds, 50 °C for 30 seconds, and 72 °C for 45 seconds, concluding with a final extension at 72 °C for 10 minutes. Post-amplification, the amplicons were quantified using the Qubit 2.0 Fluorometer with a dsDNA HS Assay Kit (Life Technologies, Grand Island, NY), and confirmed via 2% TBE agarose e-gel (Life Technologies, Grand Island, NY). The amplicons were pooled at equimolar concentration using the liquid handler (Opentrons® OT-2), and purified using AMPure SPRI beads (Beckman Coulter, Brea, CA). The final amplicons library quality and quantity were assessed with the Bioanalyzer DNA 1000 Kit (Agilent, Santa Clara, CA) and the Qubit 2.0 Fluorometer with a dsDNA HS Assay Kit, respectively. The final pooled amplicons library was diluted to 2 nM, PhiX spike-in control was added at a 35% final concentration, and the samples were sequenced on the Illumina NextSeq500 Platform producing paired-end 151 x 151 bp reads. Primer sequences and adapters were trimmed using the cutadapt plugin in QIIME2.^44^ DNA sequences underwent quality filtering, denoising, and chimeral filtering using DADA2.^45^ Taxonomy was assigned to amplicon sequence variants (ASVs) using the SILVA v138 database.^46^ Qiime artifacts were read into R for analysis using the qiime2r package (v0.99.6).

### Flow cytometry of lung cells

A single cell suspension was collected and immediately stored in phosphate-buffered saline, supplemented with 5 mM ethylenediaminetetraacetic acid, and 2% (v/v) heat-inactivated fetal bovine serum. 500,000 to 1,000,000 cells were washed once in this mixture and then analyzed by flow cytometry. The following stains were used: LIVE/DEAD™ Fixable Aqua Dead Cell Stain Kit (Thermo Fisher, Waltham, MA), APC-conjugated anti-mouse CD45 (Clone 30-F11, BD Biosciences, Franklin Lakes, NJ), BV786-conjucated anti-mouse CD326 (Clone G8.8, BD Biosciences, Franklin Lakes, NJ), BV421-conjucated anti-mouse CD31 (Clone 390, BD Biosciences, Franklin Lakes, NJ), BV605-conjugated anti-mouse CD19 (Clone 1D3, BD Biosciences, Franklin Lakes, NJ), PerCP-conjugated anti-mouse CD3e (Clone 145-2C11, BD Pharmigen, Franklin Lakes, NJ), and either the MitoProbe™ TMRM Assay Kit for Flow Cytometry (Thermo Scientific, Waltham, MA) or the MitoTracker^TM^ Dye for Flow Cytometry (Thermo Scientific, Waltham, MA). Single stain controls were used for compensation. Cell population gating is shown in Extended Data. Normalized TMRE results summarizing two independent experiments were calculated by dividing the individual mean fluorescence intensity (MFI) measurement of TMRE expression in Live CD45-CD31+CD326+ cells for each mouse in an experiment to the mean MFI of this cell type for all mice within an individual experiment; this value was then used to combine the results of two independent experiments for display in the manuscript.

For levels of splenic and T cell populations, spleen and lung tissues were isolated with single cell suspensions created of both. Cells were treated with PMA (50ng/ml) and ionomycin (1000ng/ml) for 4 hours and stained for live cells using the following stains: LIVE/DEAD™ Fixable Aqua Dead Cell Stain Kit (Thermo Fisher, Waltham, MA), PE-conjugated anti-mouse CD45 (Clone 30-F11, Fisher Scientific, Waltham, MA), FITC-conjugated anti-mouse CD3 (Clone 17a2, Fisher Scientific, Waltham, MA), BV786-conjugated anti-mouse CD4 (Clone GK1.5, Fisher Scientific, Waltham, MA), PE-Dazzle-conjugated anti-mouse-TCRβ (Clone: H57-597, BioLegend, San Diego, CA), BV421-conjugated anti-mouse IFNγ (Clone XMG1.2, BD Biosciences, Franklin Lakes, NJ), and PE-Cy7-conjugated anti-mouse IL-17A (Clone ebio17b7, Fisher Scientific, Waltham, MA). All cells were gated on live singlet populations. Hematopoietic cells were quantified as a total of CD45+ cells. T helper (Th) populations were gated on Live singlets that were CD45+CD4+TCRβ+ and represented groups of T cells producing IL-17A for Th17 cells or IFNγ in the case of Th1 cells. Cell gating strategies are shown in Extended Data.

### Untargeted metabolomic assessment of conditioned media

Conditioned media was prepared by growing *B. adolescentis* BD1 in BHI media for 24 hours, removing the supernatant, and then freezing samples at -80°C. Media samples were then reconstituted in 1:1 weight by volume acetonitrile: methanol with 1% H_2_O, agitated for 30 minutes. The aqueous layer was separated and analyzed on a SCIEX 6600+ TripleTOF Liquid Chromatography Mass Spectrometer.

### Untargeted metabolomics analysis

Analysis was done both on raw unannotated features and feature assignments with annotations from MSDIAL (v4.9.221218). In the case of annotated features, analysis was done on features present with MSDIAL fill values of > 0.95 where fill values indicate the proportion of samples where a peak was detected with high confidence. Zero replacement was conducted using a pseudocount of 0.5 and data was transformed using log_10_ transformation. Annotated metabolites with a fill > 0.95 fell into 627 different ontology groupings; these groupings were changed to large group assignments manually (Amino Acids and Derivatives, Lipids, Unknown, etc) to simplify visual display.

### Assembly and analysis of *B. adolescentis* BD1 genome

*B. adolescentis* BD1 was cultured at 37°C in an anaerobic chamber (Coy Laboratory Products) (5% H_2_, 20% CO_2_, balanced with N_2_), with DNA isolated from the overnight culture using the Qiagen High Molecular Weight DNA Kit (Germantown, MD). Samples were lysed with proteinase K, placed in buffer MB, and separated using a magnetic rack. Samples were then washed with buffer MW1 and buffer PE twice and eluted with 300 μL buffer EB. DNA fragment integrity and concentration were confirmed using the 4200 Agilent TapeStation system (Santa Clara, CA). We used a hybrid sequencing approach as described previously^47^. Briefly, for large DNA fragments, the Oxford Nanopore PCR Free Ligation Sequencing Kit (LSK109, Oxford Nanopore Technologies, Oxford, UK) was used with sequencing subsequently conducted on an Oxford Nanopore MinION sequencer (Oxford Nanopore Technologies, Oxford, UK). For short DNA fragments, 2×150bp libraries were prepared using the Illumina NovaSeq Kit and run on a NovaSeq 6000 (Illumina, Foster City, CA) instrument. The genome was assembled using Unicycler.^48^ The assembled genome was uploaded to KBase (kbase.us) with the preloaded library Distilled and Refined Annotation of Metabolism v0.1.2 tool with output data displayed using ggplot2 (v3.5.1).

### Lactate quantification

*L*-lactate was measured in bacterial cell supernatants, mouse cecal contents, and mouse stool samples. *B. adolescentis* was grown as described above. In the case of stool specimens, a frozen specimen was weighed and added to 500 μL of autoclaved water and incubated at room temperature for 30 minutes. For gut samples, 50 μL of the sample was measured using the *L-*lactate assay. *L*-lactate measurements were made using the *L*-lactate Assay Kit (Cambridge, UK). Briefly, a master mix was created using the included assay buffer, lactate enzyme mix, and the oxired probe. The included lactate standard was arrayed onto a 96-well plate followed by samples in individual wells to a volume of 50 μL. The master mix was then added to each well at 50 μL. Two to three background wells were included per assay without the oxired probe. The plate was incubated for 30 minutes at room temperature, and colorimetric measurements were made using 570 nm measurements on an Agilent BioTek Microplate Reader (Santa Clara, CA).

### Bioenergetic assessment of lung epithelial cells

A lung epithelial cell line (BEAS-2B cells) was obtained from the UCSF Cell and Genome Engineering Core (cgec.ucsf.edu) and maintained in Dulbecco’s Modified Eagle’s Medium (DMEM) (Fisher Scientific, Waltham, MA) supplemented with 10%(v/v) heat-inactivated fetal bovine serum (Life Technologies, Waltham, MA) and 100 Units/mL penicillin and streptomycin (Thermo Fisher, Waltham, MA). *L*-lactate (Fisher Scientific, Waltham, MA) was solubilized directly in H_2_O and pH adjusted to 7.4 followed by 0.22 μM filtration. 40,000 BEAS-2B cells were seeded in 96-well Agilent Seahorse XF plates. One day later, cells were treated with the indicated compounds at multiple concentrations and incubated for 24 hours. Bioenergetic assessment was done using the Agilent Seahorse XF system (Santa Clara, CA) using the Cell Mito Stress Test with 1.5 μM oligomycin and 2.0 μM FCCP based on a five-point FCCP and OCR titration to ensure average starting absolute OCR measurements of 100 pmol/min and maximal respiration crossed baseline OCR measurements and plateaued. Cells were imaged for both pre- and post-bioenergetic assessment with 1:10,000 Hoechst 34580 (Thermo Fisher, Waltham, MA) in line with protocol recommendations for normalization by Agilent. Cell imaging was done on a BioTek Cytation 5 microscope with image capture and cell count analysis obtained using the Gen5 software (Santa Clara, CA). Normalization was done on a per live cell basis.

### Quantification of *B. adolescentis* in cultures and mice

*B. adolescentis* was grown *in vitro* in BHI with the overnight culture serially diluted to 10^3^, 10^4^, 10^5^,10^6^, 10^7^, 10^8^, and 10^9^ dilutions. 25 μL of each dilution was plated onto solid BHI plates with the resulting colonizes calculated. Using this method, 24 hour cultures were found to have an average of 1.8 x 10^8^ colony-forming units (CFU) per mL. DNA was extracted from a single 24-hour culture and used to create a standard curve. The ZYMO 96-MagBead DNA kit was used for DNA extraction. Briefly, cells were spun down and added to the lysing plate in 650 μL of the ZymoBIOMICS lysis solution. Samples were disrupted for 5 minutes in a Biospec beadbeater (Bartelsville, OK) and then incubated at 65°C for 10 minutes. 200 μL of supernatant was then transferred to a deep-well plate and treated with 600 μL ZymoBIOMICS MagBinding Buffer with 2-Mercaptoethanol. 25 μL of magnetic beads were then added to each well and washed with MagWash1 once and MagWash2 twice. Residual ethanol was evaporated at 65°C for 10 minutes and samples were eluted in 100 μL nuclease-free H_2_O and captured on a magnetic plate. In the case of stool pellets from GF or *B. adolescentis* colonized mice, the identical process of DNA extraction was conducted starting with 50 mg stool from each mouse.

Once DNA was obtained, samples were analyzed using quantitative polymerase chain reaction-based assessment of bacterial load. Primers were designed for *B. adolescentis* using the speciesprimer^49^ (v2.1) primer design tool. Primers targeting the *B. adolescentis* endolytic murein-transglycosylase (mltG) gene had the highest primer design score (Forward 3’-GGGAGACAAAGACACGACGT-5’; Reverse 5’-TCCGCCTTGACCAGATTTCG-3’). A 9 μL mixture of SYBR Green I (Sigma, St. Louis, MO), KAPA HiFI PCR kit (KAPA HiFi, Cape Town, South Africa), 1 μM of Forward and Reverse primers, and nuclease free H_2_O was prepared for each reaction. A volume of 1 μL of extracted DNA was added to each reaction, and a standard curve was created from a *B. adolescentis* overnight culture-confirmed to have 1.09^8^ CFU per mL. Reactions were conducted on a BioRad CFX 384 real-time PCR instrument with 5 min at 95°C, 20x (20 sec at 98°C, 15 sec at 55°C, 60 sec at 72°C), and hold at 4°C. Reaction quality was assessed by evaluating amplification curves and melting curves for all products.

### Metabolomic analysis from the <u>E</u>arly <u>A</u>ssessment of <u>R</u>enal and <u>L</u>ung Injury (EARLI) Cohort

As described in the prior publication^50^, this study was approved by the University of California San Francisco (UCSF) institutional review board (IRB 310987). 198 critically ill patients at UCSF Moffitt-Long hospital and San Francisco General Hospital were enrolled with plasma samples collected within 24 hours of ICU admission. ARDS was adjudicated using American-European Consensus Conference (AECC) definitions using two person review of chest radiographs^51^. Untargeted metabolomic profiling was conducted by Metabolon (Durham, NC) on citrated plasma using three complementary methods: ultrahigh performance liquid chromatography/tandem mass spec UHLC/MS/MS2 for basic species, UHLC/MS/MS2 for acidic species, and UHLC/MS/MS2 for lipids. Annotated metabolites were batch-normalized and missing values were imputed using the minimum value of a given metabolite across all batches by Metabolon. Imputed data was then log_10_ transformed. A linear model was calculated using the stats package in R with the lm command (Metabolite value ∼ AECC ARDS* APACHE3 score). Resulting *p*-values for the APACHE3 term were corrected using the Benjamini-Hocheberg method. Pearson’s *r* values and *p*-values shown in the main figures were calculated using the cor_test command in the rstatix package on a per metabolite basis and corrected using the Benjamini-Hochberg method where indicated. A euclidean distance matrix was generated using log_10_ transformed values using the stats package.

### Other tools for the display of figures and statistical analysis

Figures in the paper were made using R (v4.2.1) with tidyverse (v2.0.0). Statistical comparisons were completed using rstatix (v0.7.2), and ggpubr (v0.6.0). The preliminary drafts of text were drafted with spell check and grammar suggestions using Grammarly.

## RESULTS

Multiple prior studies have implicated T helper 17 (Th17) activation in ARDS^23,52,53^, prompting us to initially focus on bacteria known to impact Th17 activation^54^. We previously isolated a strain of *Bifidobacterium adolescentis* (strain BD1) from a human subject that is a potent Th17 inducer in the small intestine^38^; however, the impact of *B. adolescentis* BD1 on systemic immunity remained unknown. We colonized GF C57BL/6J adult male mice (n=3-6/group) with *B. adolescentis* BD1 for 2 weeks and then harvested gut, spleen, and lung tissue for analysis. As expected, *B. adolescentis* BD1 colonized the distal gut at high levels (8.3±2.3 x 10^8^ CFU/g wet weight; **Extended Data Fig. 1a**) but was undetectable in the lung (**Extended Data Fig. 1b**). Spleen and lung weight were unaffected by colonization (**Extended Data Figs. 1c,d**). Immune profiling by flow cytometry revealed that total CD45+ cells, T helper 1 (Th1), and Th17 cells were comparable between GF and mono-colonized mice in both organs (**Extended Data Figs. 1e-n** and **Extended Data Fig. 2**), which together with prior data^38^ indicates that the immune effects of this strain are restricted to the gastrointestinal tract.

Next, we turned to bulk RNA sequencing (RNA-seq) to assess if *B. adolescentis* colonization had an impact on the non-immune cells within the lung (**Supplemental Table 1a**). Remarkably, we found that the overall lung transcriptome was significantly altered by *B. adolescentis*, separating clearly on principal coordinate 1 (Fig. 1a). We detected a total of 2,801 differentially expressed genes (DEGs, *p*_adj_<0.1), including 1,329 up-regulated and 1,472 down-regulated in response to colonization (**Supplemental Table 2a**). These DEGs showed a non-random distribution across chromosomes, with a marked and significant up-regulation of transcripts encoded by the mitochondrial genome (Fig. 1b). Genes with known mitochondrial activity showed signs of upregulation independent of their location in the nuclear genome, including a general increase in expression level for nuclear genome-encoded mitochondrial genes in response to *B. adolescentis* (**Figs. 1c,d**). Overall *B. adolescentis* led to a significant up-regulation of 162 mitochondrial genes, including 7 in the mitochondrial genome and 155 genes with mitochondrial activity from the nuclear genome. Consistent with this broad pattern of mitochondrial gene induction, the *B. adolescentis* DEGs encompassed all five electron transport chain complexes (Fig. 1e). Similar effects on mitochondrial transcript levels were observed in colon tissue following *B. adolescentis* colonization (**Extended Data Fig. 3** and **Supplemental Tables 1b,2b**).

**Figure 1:**
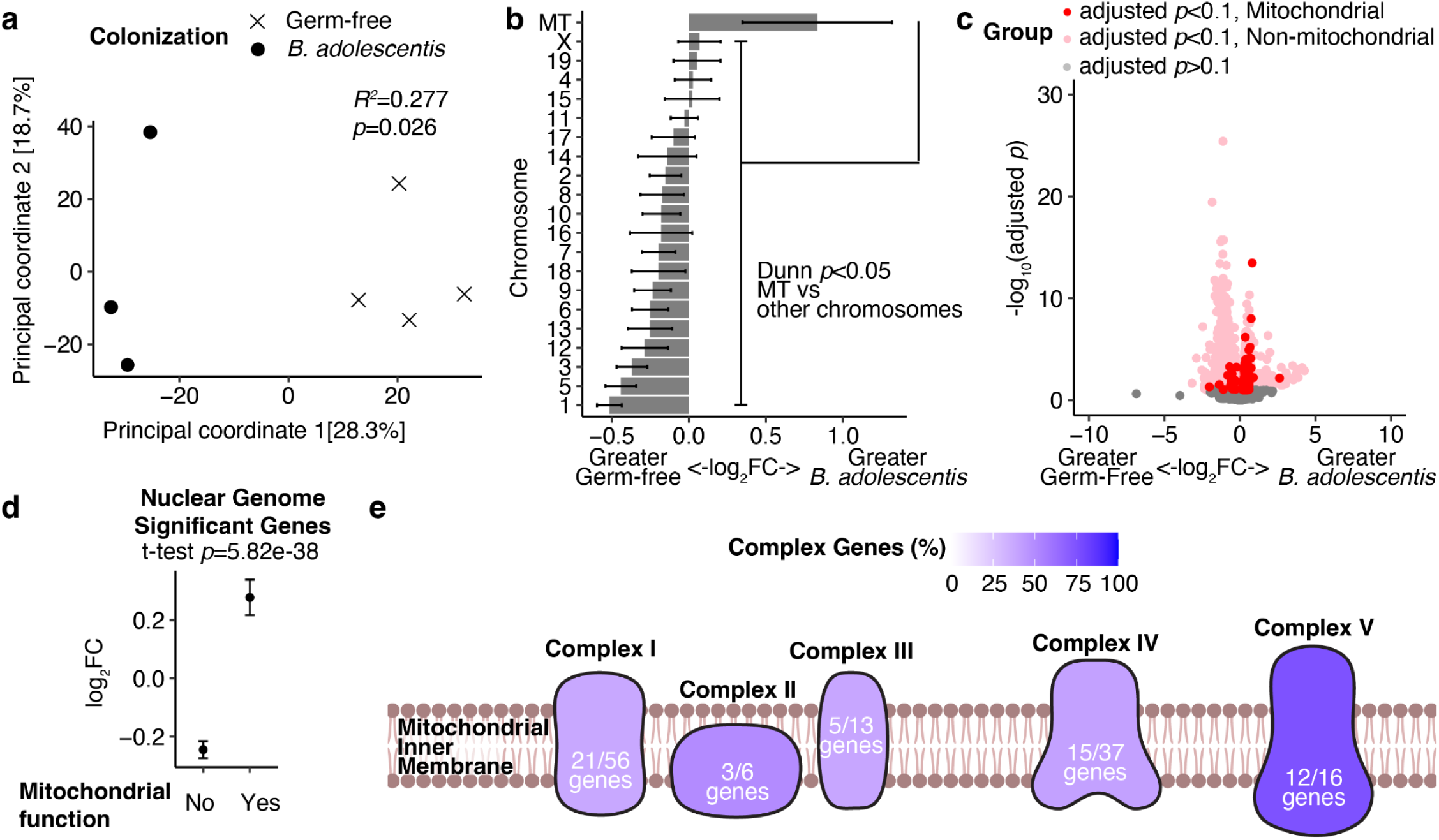
*B. adolescentis* stimulates the expression of mitochondrial electron transport genes within the mouse lung. **(a)** Principal coordinate analysis of bulk RNA sequencing from lung tissue isolated from GF or *B. adolescentis* colonized mice. PERMANOVA testing *R^2^* and *p*-value annotated between colonization groups. **(b)** Differentially expressed genes between GF and *B. adolescentis*-colonized mice plotted by chromosome of origin with mean and 95% confidence interval indicated by error bars. A Dunn’s post hoc test comparing the mitochondrial (MT) genome to all chromosomes of the nuclear genome individually and summarized is annotated on the graph. **(c)** RNA sequencing data plotting -log_10_(adjusted *p*-value) against log_2_(fold-change) between GF and *B. adolescentis* colonized groups highlighting genes with mitochondrial activity independent of chromosome of origin. **(d)** Mean and 95% confidence interval for differentially expressed genes between GF and *B. adolescentis* colonized mice comparing those with mitochondrial function to those without encoded in the nuclear genome. **(e)** *B. adolescentis* induced electron transport chain genes as a percentage of each mitochondrial electron transport complex. Significantly upregulated gene components for each mitochondrial electron transport chain upregulated by *B. adolescentis* over total detected components. n=4 GF and n=3 *B. adolescentis* colonized mice.

We performed an independent experiment to assess the reproducibility and specificity of these findings. GF C57BL/6J adult male mice (n=3-4/group) were colonized with *B. adolescentis* BD1, *Eggerthella lenta* DSM2243 (another member of the *Actinomycetota* phylum), or a complex mouse gut microbiota for 2 weeks. We then harvested lung tissue for bulk RNA-seq analysis (**Supplemental Table 1c**). We compared each colonization group to GF controls (**Figs. 2a-c**). Consistent with our original experiment, *B. adolescentis* altered the overall lung transcriptome (**Figs. 2a**) leading to a significant up-regulation of 957 genes, including 34 mitochondrial transcripts (**Figs. 2d,e** and **Supplemental Table 2c**). The conventionalized (CONV-D) mice colonized with a complex mouse gut microbiota showed a consistent impact on lung mitochondrial gene expression to *B. adolescentis* colonized mice, exhibiting a significant shift in overall expression relative to GF controls (Fig. 2b) and a significant up-regulation of mitochondrial genes (**Figs. 2f,g** and **Supplemental Table 2c**). In contrast, *E. lenta* had a more modest impact on the lung transcriptome (Fig. 2c) with 138 DEGs (95 up- and 43 down-regulated) and no detectable enrichment for mitochondrial transcripts (**Figs. 2h,i** and **Supplemental Table 2c**).

**Figure 2:**
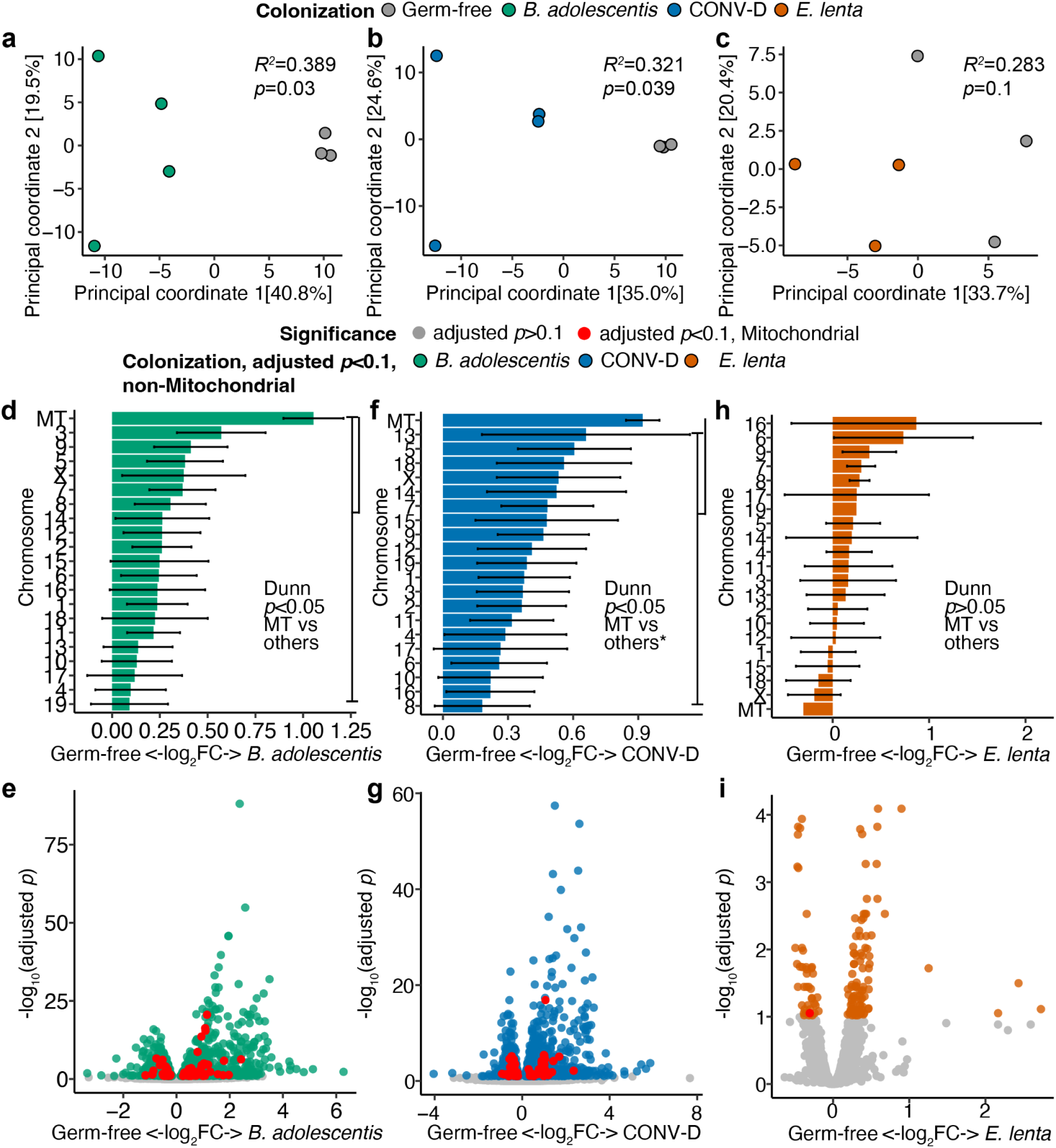
The gut microbiota induces mitochondrial gene expression in lung tissue. **(a-c)** Principal coordinate analysis of bulk lung RNA-seq datasets comparing each group relative to GF mice. The legend for panels **d-i** is above panels **d,f,h. (d, f, h)** Differentially expressed genes between GF mice and mice of each colonization state are plotted by chromosome of origin with mean and 95% confidence interval indicated by error bars. A Dunn’s post hoc test comparing the mitochondrial (MT) chromosome to all others individually and summarized is annotated on the graph. The asterisk indicates for the conventionalized (CONV-D) group *p*=0.056 for the mitochondrial and X chromosomes. **(e, g, i)** The -log_10_(adjusted *p*-value) is plotted against log_2_(fold change) (log_2_FC) between GF mice and the indicated colonized groups highlighting genes with mitochondrial activity independent of the chromosome of origin. n=3 GF, n=4 CONV-D, n=4 *B. adolescentis* mice, and n=3 *E. lenta* mice for all panels.

Comparing all colonization states directly, the CONV-D and *B. adolescentis* groups clustered separately from the GF and *E. lenta* colonized mice (**Extended** Fig. 4a). *B. adolescentis* and CONV-D had the same effect on 845 genes (**Extended** Fig. 4b), including genes spanning four mitochondrial electron transport chain complexes (**Extended** Fig. 4c). All CONV-D mice were colonized by a member of the *Bifidobacterium* genus (3.6±1.1% relative abundance), including a strain of *B. pseudolongum* species (**Extended** Fig. 4d). However, the human-associated species^55,56^ *B. adolescentis* was undetected despite deep 16S rRNA gene sequencing (4.46±0.35 x 10^8^ sequencing reads per sample; **Supplemental Table 1d**). Taken together, these results suggest that the observed impact on mitochondrial gene expression generalizes to other bifidobacteria but not more distantly related members of the *Actinomycetota* phylum.

Next, we used single-cell RNA sequencing (scRNA-seq) to determine if the bulk tissue up-regulation of mitochondrial gene expression was driven by a change in cell type. We colonized a mouse with *B. adolescentis* for 2 weeks and separated it from its GF littermates. We sorted live CD45- and CD45+ cells from lung tissue and immediately prepared a scRNA-seq library. We sequenced a range of 1.38 - 5.64 x 10^3^ cells per sample, obtaining 3.04±0.50 x 10^8^ reads per sample (**Supplemental Table 1e**). Cells were annotated with the Mouse Cell Atlas^42,57^. Consistent with prior studies^58,59^, the CD45-cells were primarily composed of endothelial, epithelial and stromal cells (Fig. 3a and **Extended Data Fig. 5a**), whereas the CD45+ cells were annotated as macrophages, T cells, B cells, and other hematopoietic cell types established to be present in the lung and associated vascular compartments (**Extended Data Figs. 5b-c**). The cellular composition of both CD45-(Fig. 3b) and CD45+ (**Extended Data Fig. 5d**) cells were highly correlated between mice (*r^2^*>0.90). Flow cytometry-based analysis confirmed that the proportion of CD45+ cells was stable with respect to colonization state (**Extended Data Figs. 1e-f**). Thus, our bulk RNA-seq data was likely driven by cell intrinsic shifts in gene expression, not an overall change in cell type or lung immune infiltration.

**Figure 3:**
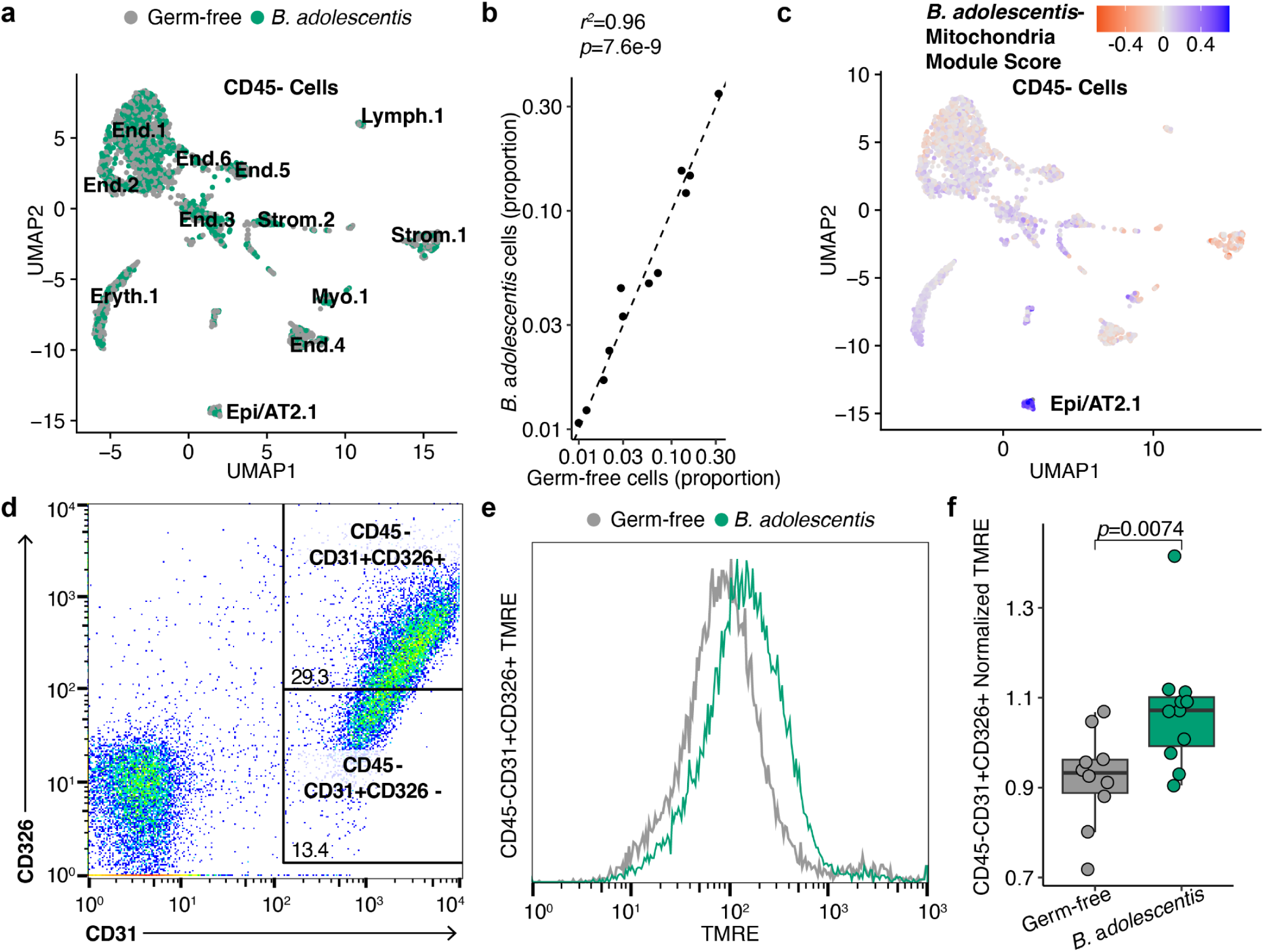
*B. adolescentis* stimulates mitochondrial activity in alveolar epithelial cells. **(a-c)** scRNA-seq of live CD45-cells from a single GF or *B. adolescentis* colonized mouse. **(a)** Cluster assignments completed with Seurat and annotated with the Mouse Cell Atlas. Individual cells are colored by colonization state. **(b)** The proportion of each cluster for *B. adolescentis* colonized vs GF mice is plotted. The Pearson’s correlation *r^2^* and *p*-value are annotated above the plot. **(c)** A module score was calculated in Seurat from mitochondrial genes differentially expressed by *B. adolescentis* colonization (adjusted *p*<0.1), and individual cells are colored by module score. **(d)** Flow cytometry for CD326 high cells enriched for alveolar epithelial cells. **(e)** Mean fluorescence intensity (MFI) of tetramethylrhodamine methyl ester (TMRE) signal for CD45-CD31+CD326+ cells from **(d)**. **(f)** TMRE levels in CD45-CD326+CD31+ cells normalized to the mean MFI of all mice in an individual experiment. n=10-11 mice per group in **(d-f)** from two different experiments.

To identify which cell type was most responsive to colonization, we calculated a module score of mitochondrial DEGs within the scRNA-seq data. DEGs were calculated between clusters in CD45- or CD45+ cells, and DEGs were then filtered for mitochondrial genes (Fig. 3c and **Extended** Fig. 5e). *B. adolescentis*-induced mitochondrial genes were highly expressed in the subset of non-hematopoietic cells, annotated as epithelial/alveolar type 2 cells (Fig. 3c). Cells in this cluster had a mean z-score value of 4.14 compared to the cluster average of 0.03±1.37 for all CD45-cells. This is notable because epithelial cells make up the majority of the surface area of the lung^1^, and alveolar type 2 cells play key roles in surfactant production and homeostasis^60^. In contrast, none of the CD45+ cell types showed significant variation in elevated levels of mitochondrial gene expression based on *B. adolescentis* colonization (-0.34±0.93 cluster average, no cluster with |mean Z score| > 4; **Extended** Fig. 5e).

**Figure 4:**
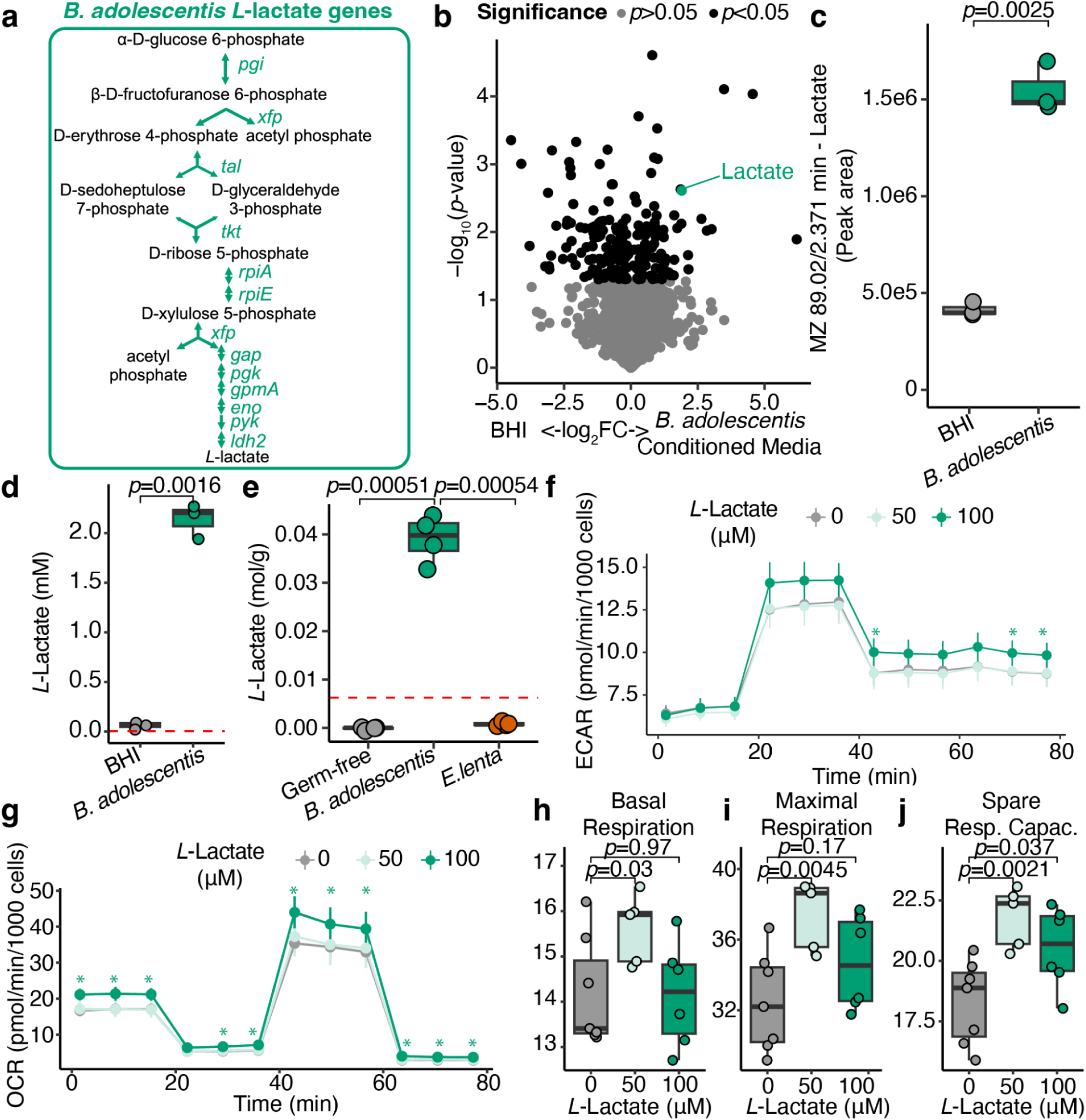
The bifidobacterial metabolite *L*-lactate is elevated in mice and increases mitochondrial efficiency in pulmonary epithelial cells. **(a)** The complete pathway for *L–*Lactate production from the *B. adoelscentis* BD1 genome. *B. adolescentis* BD1 encodes genes required for all chemical transformations. Gene symbols shown in green italics. Select intermediate and end products are displayed in black. **(b)** Liquid chromatography-mass spectrometry-based metabolomic profiling of *B. adolescentis* conditioned media plotting -log_10_(*p*-value) against log_2_(fold change) (log_2_FC). Predicted annotations were conducted by MSDIAL, and the negative ionization feature MZ 89.02, Retention time 2.371 minute predicted to be Lactate is annotated and colored in green. n=3 per group. **(c)** The lactate feature (MZ 89.02, Retention time 2.371 minutes) peak area is plotted for BHI or *B. adolescentis* conditioned media. (d) *L*-lactate production by enzymatic assay for *B. adolescentis* grown *in vitro* for 24 hours (n=3 biological replicates per group). (e) *L*-lactate levels in cecal contents from GF, *B. adolescentis* colonized, or *E. lenta* colonized mice (n=4 mice per group). **(c-e)** The red dashed line indicates the detection limit. The *p*-values indicate *t-*tests between the indicated groups. **(f-j)** Bioenergetic profiling of BEAS-2B cells pretreated with *L*-lactate at two concentrations. **(e)** Extracellular acidification rate (ECAR) normalized to 1000 cells or **(f)** Oxygen consumption rate (OCR) normalized to 1000 cells is plotted against time. Asterisks in **(f-g)** indicate Wilcox *p*<0.05 for the concentration of the indicated compound versus the 0 μM control group for a given timepoint. **(h)** Basal respiration, **(i)** Maximal Respiration, **(j)** Spare Respiratory Capacity for *L*-lactate treated BEAS-2B cells. **(f-j)** Each experiment had 4-8 replicates per group in an experimental plate, with 2-4 replicates done for each compound.

**Figure 5:**
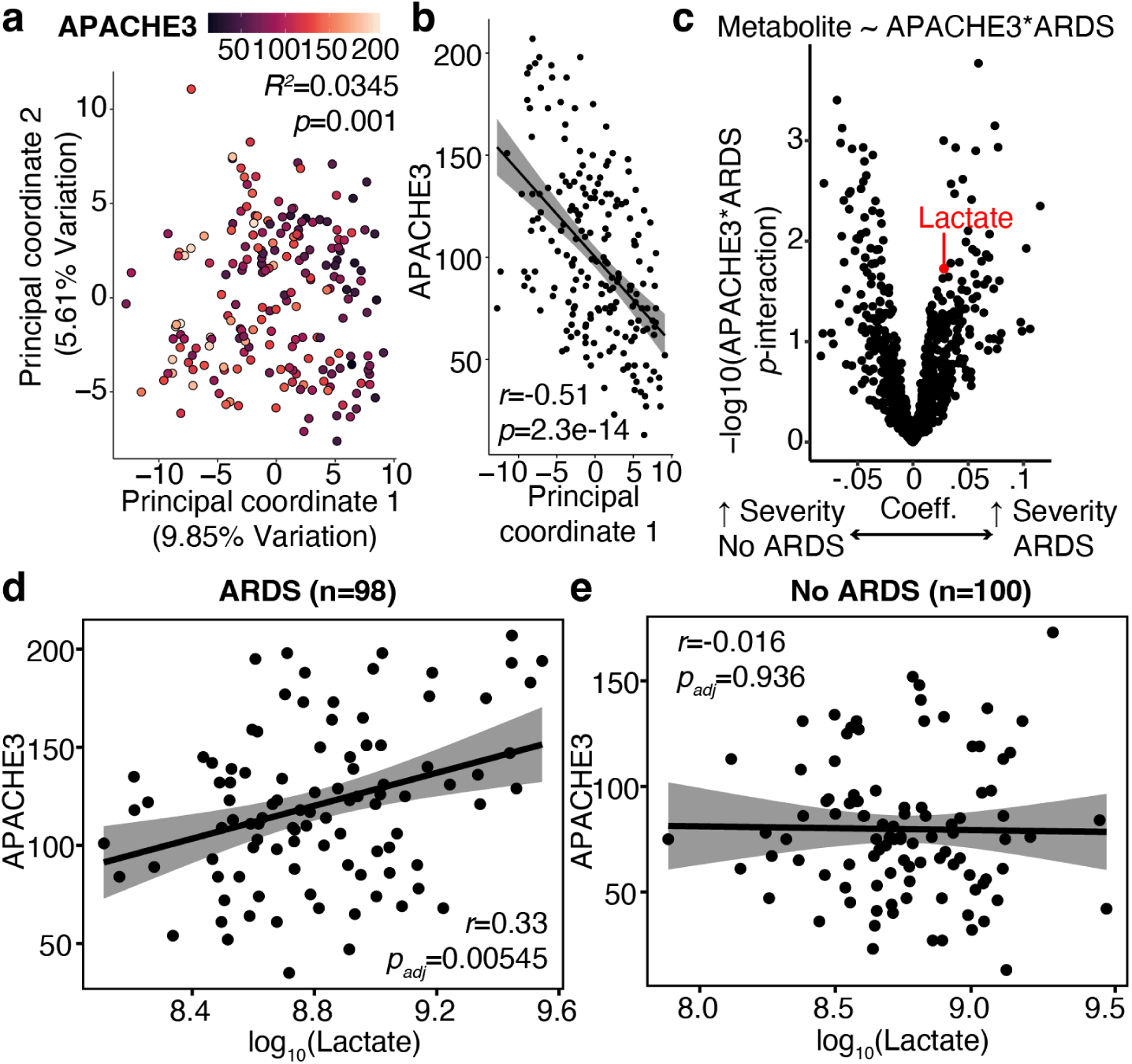
Serum lactate correlates with severity of illness in critically ill patients with ARDS. **(a)** Principal coordinate analysis displaying the two largest components of variation for a Euclidean distance matrix generated from 970 serum metabolites. PERMANOVA testing results for Acute Physiology and Chronic Health Evaluation (APACHE) 3 scores are annotated above the figure. **(b)** APACHE3 score versus Principal coordinate 1 from panel **(a)** with Pearson’s *r* and *p*-value annotated. **(c)** A linear model was created with the equation annotated above the graph with the acute respiratory distress syndrome (ARDS) *APACHE3 interaction coefficient plotted against its *p*-value. **(d-e)** APACHE3 scores plotted against lactate measurements for patients **(d)** with or **(e)** without ARDS with Pearson’s correlation coefficient annotated along with *p*-value adjusted for multiple testing. n=100 patients with early sepsis without ARDS and n=98 patients with early sepsis and ARDS.

Taken together, our bulk and scRNA-seq results show that *B. adolescentis* colonization of the gut induces mitochondrial gene expression in the lung, most notably in epithelial cells. However, the functional consequences for mitochondrial activity remained unclear. We conducted flow cytometry of freshly isolated lung cells to assess epithelial cell mitochondrial membrane potential and mitochondrial biomass (**Extended Data Fig. 6**, **Extended Data Fig. 7** and Fig. 3d). Mitochondrial biomass in lung epithelial cells was unaffected by gut bacterial colonization (**Extended Data Fig. 7**). However, mitochondrial membrane potential was significantly increased in response to colonization with *B. adolescentis* (**Figs. 3d-f**). Maintenance of membrane potential depends on the mitochondrial-cytoplasmic proton gradient with abnormally increased potential being indicative of elevated levels of reactive oxygen species^61^.

Next, we sought to better understand the mechanism through which a bacterium within the gut could influence lung gene expression and mitochondrial activity. We hypothesized that a secreted metabolite could be signaling directly to lung epithelial cells based upon the following observations: (*i*) the overall cell composition of lungs was unaffected by colonization in our scRNA-seq data (**Figs. 3a,b**); (*ii*) the level of migratory immune cells was unaffected by flow cytometry (**Extended Data Fig. 1** and **Extended Data Figs. 5b-d**); and (*iii*) mitochondria are known to be impacted by a variety of gut bacterial metabolites^62^.

To identify potential secreted metabolites, we assembled a high-quality genome of *B. adolescentis* BD1 using hybrid long- and short-read sequencing (**Supplemental Table 3a**). A total of 1,803 coding sequences were found across the 2.23 Mbp genome in a single contig (**Supplemental Table 3b**). We assessed the metabolic capacity of the *B. adolescentis* strain BD1 genome. This analysis revealed 8 complete genetic pathways in carbohydrate and short-chain fatty acid metabolism (**Extended Data Fig. 8a**). As expected, this included a complete pathway for the fermentation of *L-*lactate (Fig. 4a and **Extended Data Fig. 9**), which is a common end-product of all bifidobacteria^63^. We used untargeted metabolomics to validate the computationally predicted metabolites of *B. adolescentis*, which revealed 1,391/2,805 features were significantly enriched following *in vitro* culture in rich media (**Extended Data Fig. 8b** and Fig. 4b), including lactate (**Figs. 4b-c**). We further validated the production of *L*-lactate by *B. adolescentis* during *in vitro* growth using an isomer-specific enzymatic assay (Fig. 4d). Using this assay, we also detected a significant increase of *L*-lactate within the cecum of *B. adolescentis* mono-colonized mice relative to GF or *E. lenta* mono-colonized controls (Fig. 4e).

*L*-lactate was sufficient to alter the mitochondrial activity of pulmonary epithelial cells. We utilized the bronchial epithelial cells transformed by adenovirus-12 SV40 (BEAS)-2B cell line^64^ to perform live cell imaging and bioenergetic assessment following a 24-hour exposure to *L*-lactate (**Extended Data Fig. 10**). Administration of 100 µM *L*-lactate adjusted to pH 7.4 had a modest impact on extracellular acidification rate (ECAR; Fig. 4f) and resulted in significant elevation in oxygen consumption (OCR; Fig. 4g). This effect was dose-dependent, with no significant impact of 50 µM pH-adjusted *L*-lactate on the overall OCR/ECAR curves relative to vehicle controls (**Figs. 4f,g**). All three summary statistics were also significantly impacted by pH-adjusted *L*-lactate at variable concentrations, including elevated basal respiration (Fig. 4h), maximal respiration (Fig. 4i), and mitochondrial spare respiratory capacity (Fig. 4j).

Finally, we sought to assess the potential translational relevance of *L-*lactate for ARDS patients, given *L-*lactate is an established biomarker of critical illness^65^. We re-analyzed published serum metabolomic profiles of 198 patients enrolled in the <u>E</u>arly <u>A</u>ssessment of <u>R</u>enal and <u>L</u>ung Injury (EARLI) intensive care unit patient cohort^37^. This subset of the EARLI cohort included comparable numbers of patients with and without ARDS (98 ARDS, 100 no ARDS). Principal coordinates analysis revealed a clear separation of serum metabolomes based on illness severity (Fig. 5a) that was statistically significant by PERMANOVA (*R^2^*=0.0345, *p*-value=0.001). Principal coordinate 1 was also significantly associated with illness severity (Fig. 5b).

To identify individual metabolites of interest, we evaluated metabolite abundance as a function of a multiple-component linear regression model based on disease severity and ARDS status, including an interaction term that identifies differences in metabolite association between patients with and without ARDS (Fig. 5c). This analysis revealed 49 ARDS-associated metabolites, including 16 that were associated with greater illness severity and 33 that were associated with lower illness severity in patients with ARDS (**Supplemental Table 4**). Notably, this included lactate and 24 fatty acids (Fig. 5c and **Supplemental Table 4**) linked to mitochondrial activity^66^. Fourteen of these fatty acids were carnitine derivatives, which is significant because of the role of bacteria in both carnitine anabolism and catabolism^67,68^. Lactate was significantly associated with illness severity in patients with ARDS (Fig. 5d) but was unrelated to illness severity in patients without ARDS (Fig. 5e).

## DISCUSSION

Although there is an established role for the gut microbiome in educating the immune system and therein affecting the hematopoetic microenvironment of the lung, many open questions remain regarding whether or how microbiota-derived metabolite signals to distant organs affect non-hematopoetic cell types within lung tissue^69–72^. While the impact of the microbiome on inflammatory cytokines or cell types is important including in ARDS^31,33^, we lack a fundamental understanding of how high biomass gut isolates may impact the non-migratory cell types within the lung prior to injury, which is a pivotal starting point for many lung diseases like ARDS. Our study shows the lung is highly responsive to colonization with high biomass gastrointestinal isolates and that lung epithelial mitochondria are stimulated in response to colonization with a complete gut microbiota. This remote interaction is mediated by the bifidobacterial metabolite *L*-lactate, though other bacterial metabolites may be capable of producing similar responses and more complex connections between a complete gut microbial community and the lung potentially exist. While *L*-lactate was sufficient to increase the mitochondrial activity of lung epithelial cells, in excess it was correlated with increased disease severity in patients with ARDS where lung epithelial injury and associated mitochondrial dysfunction have been previously established ^2,20,21^.

These findings underpin the importance of better defining the microbial contributions to ARDS. While the role of bacteria as inciting causes of inflammation has been previously investigated^34^, we show a clear role for gut-derived bacterial metabolites in altering the metabolism of pulmonary epithelial cells. A prevailing interpretation of the role of the gut microbiome in ARDS has been on the impact of gut isolates on immune development^31,33,73^. A connection between the gut microbiota and mitochondrial metabolism adds to this work and shows that like gastrointestinal epithelial cells^74–76^, pulmonary epithelial cells are in communication with high biomass gastrointestinal isolates. Recently, the microbiome has been suggested to impact redox balance in host tissues^32^, and our data complements these observations in the lung because mitochondria maintain cellular redox potential^77^. Furthermore, serum mitochondrial nucleic acid has been a predictor of ARDS disease severity^21^, and our data suggests microbial metabolites themselves might promote ARDS by augmenting mitochondria in epithelial cells.

Mitochondria regulate host metabolism by tightly regulating both serum and cellular *L*-lactate and pyruvate levels, with lactate being a critical substrate for cellular electron transfer^78^. Our data show that *B. adolescentis* drives mitochondrial membrane potential *in vivo*, and the bifidobacterial metabolite *L*-lactate directly increases mitochondrial respiration in epithelial cells. In excess, driving this membrane potential may be a contributor to greater disease severity observed specifically in patients with ARDS. Furthermore, our observations go hand-in-hand with prior work showing a glycolytic shift in the hyperinflammatory subtype of ARDS, which is also correlated with increased mortality^20,79^.

Our study draws attention to a greater body of literature showing bidirectional interactions between mitochondria and the microbiome. Mice harboring deleterious mutations in mitochondria have distinct gut microbial communities, with a putative role for reactive oxygen species in mediating this effect^80^. The mitochondrion itself is an organelle with close genetic similarity to free-living ⍺-Proteobacteria, with hypothesized origins in a divergent proteobacterial ancestor^81^. While other groups have shown the key importance of gut microorganisms in regulating intestinal epithelial bioenergetics^75,76,82^, our own observations widen this to the responsiveness of host mitochondria to distal vascular beds. Interspecies bacterial communication occurs through autoinducers that facilitate quorum sensing^83^ and some host-derived compounds can bind to quorum sensing receptors in bacteria^84–86^. We broaden the scope of this view by showing this interaction occurs in reverse, with bacterial metabolites directly impacting host mitochondria in organs in distal anatomical compartments.

There are certainly many future avenues for investigation in this broader framework. While we show excess *L-*lactate correlates with worse disease severity in ARDS, a key issue to resolve is that mitochondrial stimulation may actually be beneficial to pulmonary epithelia in other scenarios, and the deleterious effect of *L*-lactate may be context-dependent. This aligns with other open questions regarding the role gut bacterial-induced stimulation of the mitochondria plays within ARDS. While we demonstrate that *B. adolescentis* increases mitochondrial membrane potential in lung epithelial cells, it is likely the dysregulated and excessive production of mitochondrial-derived free radicals originating within the mitochondria seem the most plausible contributors to ARDS and may be the molecular drivers of lung injury. Finally, while we focus on the role of the bifidobacterial metabolite *L*-lactate, in the host, multiple gut bacterial metabolites act on the epithelial cell simultaneously, which leaves open the possibility of synergy or antagonistic effects of different gut bacterial metabolites.

Collectively, our study shows that gut isolates make a key contribution to lung epithelial metabolism through the secretion of bioactive gut metabolites. Using an unbiased approach, we observe that lung epithelial mitochondria respond to high biomass gut isolates, and in excess, *L*-lactate is associated with greater illness severity in human subjects with epithelial injury where mitochondrial dysfunction has been established. These findings provide a new framework for understanding the impact of the microbiome on lung health and establish a role for high biomass gut isolates in driving lung cellular respiration.

## ACKNOWLEDGEMENTS

Gnotobiotic experiments were performed at the UCSF Gnotobiotics Core Facility (gnotobiotics.ucsf.edu). Sequencing was performed at Chan-Zuckerberg Biohub-San Francisco, the UCSF Genomics CoLab, and the UCSF Microbial Genomics CoLab. Metabolomics was conducted at the Quantitative Metabolite Analysis Center at UCSF (microbiome.ucsf.edu/quantitative-metabolite-analysis-center-colab-plug).

## AUTHOR CONTRIBUTIONS

Conceptualization: VU, PJT

Methodology: VU, EFO, GK, KT, RR, LRH, CC, NAL, WCC, MM, JVL, IJ, PJT

Investigation: VU, EFO, GK, KT, RTS, MA, RRR, LRH, WCC, NAL, CC, IJ, MM, JVL

Visualization: VU, GK, EFO

Funding acquisition: VU, PJT

Project administration: PJT

Supervision: PJT, AG, MM, CC

Writing – original draft: VU, PJT

Writing – review & editing: All authors

## DATA AND MATERIALS AVAILABILITY

This study did not generate new reagents. All code and data for generating figures is on GitHub at https://github.com/turnbaughlab/2025_BifidobacteriumALI. The accession number for the sequencing data reported in this paper is NCBI Sequence Read Archive BioProject PRJNA759870 which includes a full length sequence of the *B. adolescentis* genome.

## EXTENDED DATA FIGURES AND FIGURE LEGENDS

**Extended Data Figure 1:**
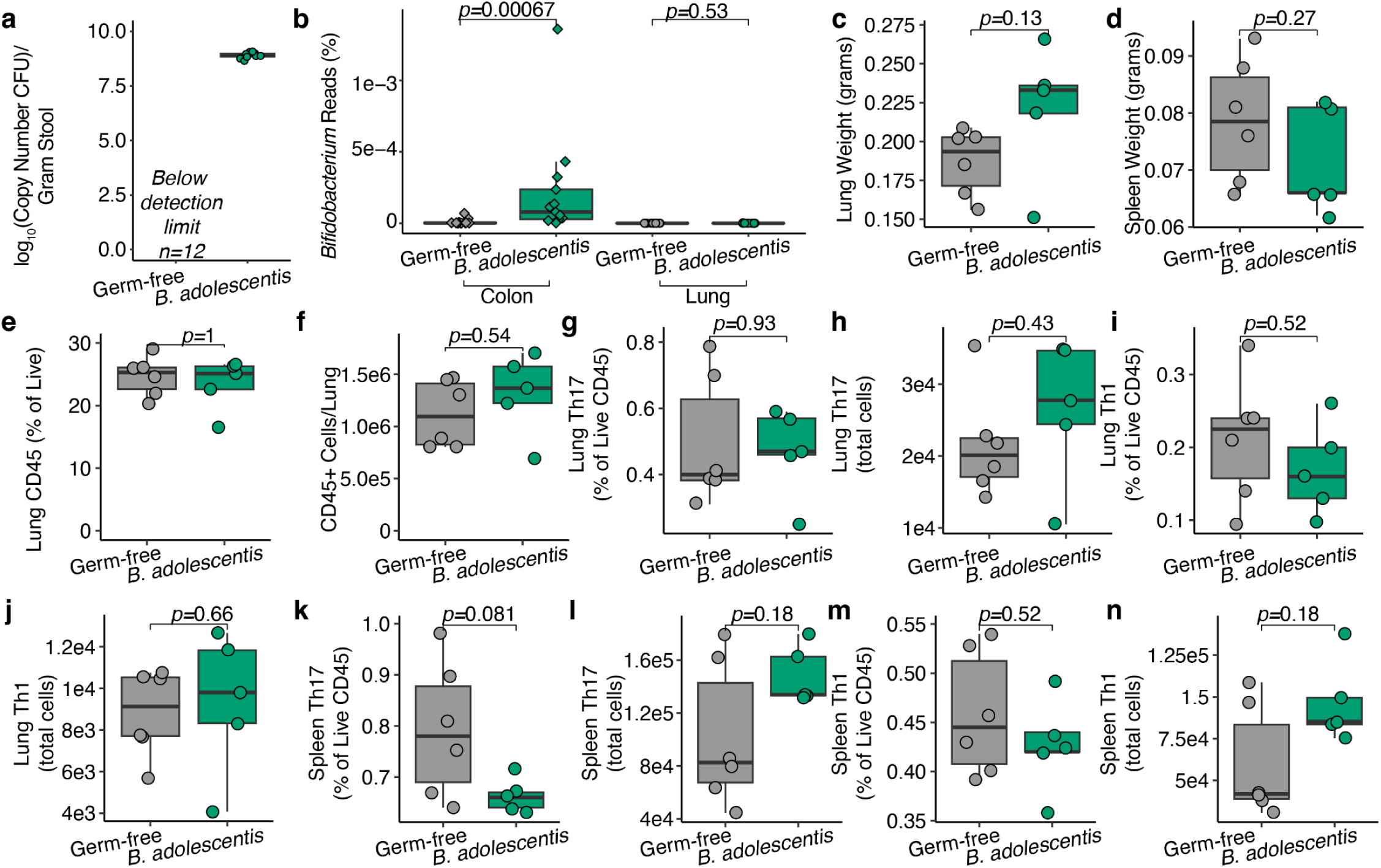
*B. adolescentis* robustly colonizes the gastrointestinal tract. **(a)** Copy number of colony-forming units (CFU) per gram of stool from GF and *B. adolescentis* colonized mice. Limit of detection 10^6.9^ (n=10 *B. adolescentis*, n=12 GF mice). **(b)** RNA sequencing reads from GF and *B. adolescentis* colonized mice from the colon or lung were input into Kraken2 using k=35, ℓ=31, and s=7 with nucleotide databases^87^. The percentage of input reads aligned with *Bifidobacterium* genera is plotted as a percentage of all reads (n=12-14 mice/group). The tissue type is indicated by the brackets below the figure, with diamonds representing colon and circles representing lung. **(c)** Total lung or **(d)** spleen weight of GF and *B. adolescentis* colonized mice. Lung CD45+ cells as a **(e)** percentage of live cells or **(f)** total number of CD45+ cells per lung. Lung Th17 cells (CD4+TCRβ+IL-17a+) as a **(g)** percentage of Live CD45+ cells or **(h)** total number of Th17 cells per lung. Lung Th1 cells (CD4+TCRβ+IFNy+) as a **(i)** percentage of Live CD45+ cells or **(j)** total number of Th1 cells per lung. Splenic Th17 cells as a **(k)** percentage of Live CD45+ cells or **(l)** total number of Th17 cells per spleen. Splenic Th1 cells as a **(m)** percentage of Live CD45+ cells or (**n)** total number of Th1 cells per spleen. n=4-7 mice per group in panels **c-n**.

**Extended Data Figure 2:**
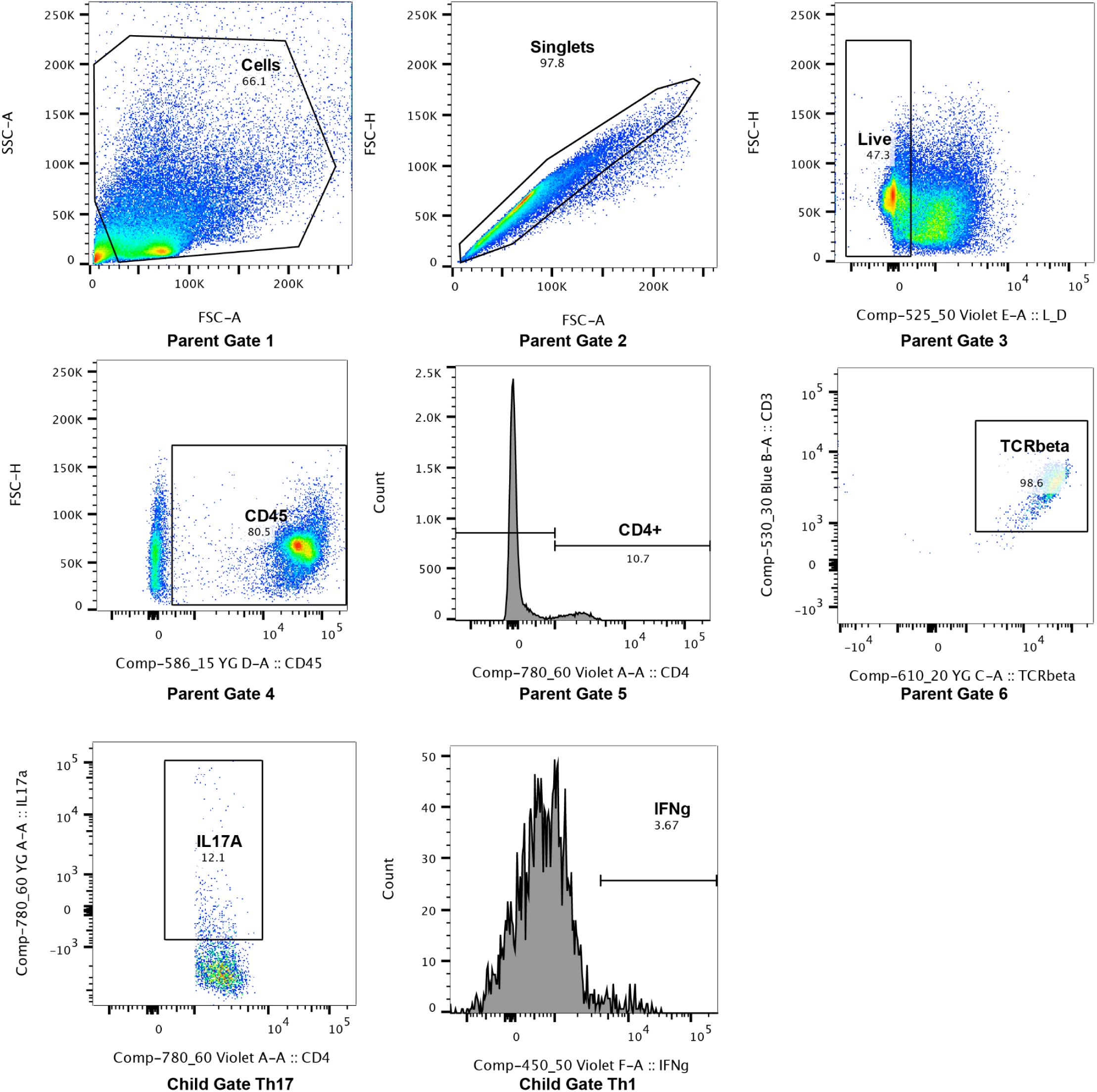
Gating strategy for flow cytometry evaluating Th1 and Th17 cells in lung and spleen. Parent gates were set on Cells (Parent Gate 1), followed by Singlets (Parent Gate 2) followed by Live Cells (Parent Gate 3), followed by CD45+ Cells (Parent Gate 4), followed by CD4+ (Parent Gate 5), and confirmed to uniformly express TCRβ (Parent Gate 6). Th17 and Th1 cells were gated on IL-17A (Child Gate Th17) or IFNγ production (Child Gate Th1).

**Extended Data Figure 3:**
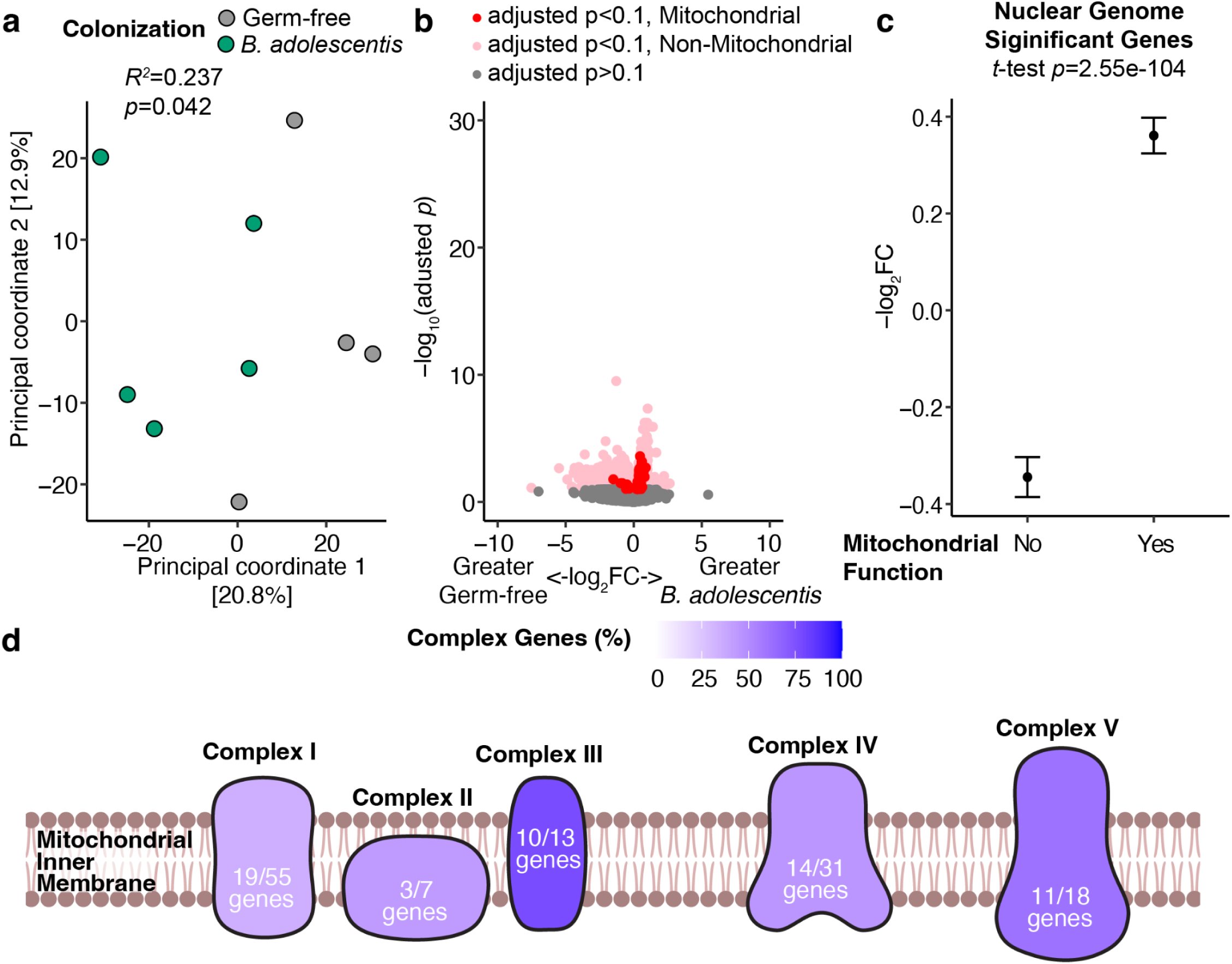
*Bifidobacterium adolescentis* induces expression of mitochondrial electron transport chain within the colon. **(a)** Principal coordinate analysis of bulk RNA sequencing from colon tissue isolated from GF or *B. adolescentis* colonized mice. PERMANOVA testing *R^2^* and *p*-value annotated between GF and *B. adolescentis* colonized groups. **(b)** RNA sequencing data plotting -log_10_(adjusted *p*-value) against log_2_(fold change) (log_2_FC) between GF and *B. adolescentis* groups highlighting genes with mitochondrial activity. **(c)** Mean and 95% confidence interval for differentially expressed genes between *B. adolescentis* colonized and GF groups comparing those with mitochondrial function to those without. **(d)** *B. adolescentis* induced electron transport chain genes as a percentage of each mitochondrial electron transport complex. Significantly upregulated gene components for each mitochondrial electron transport chain upregulated by *B. adolescentis* over total detected components. n=4 GF and n=3 *B. adolescentis* colonized mice.

**Extended Data Figure 4:**
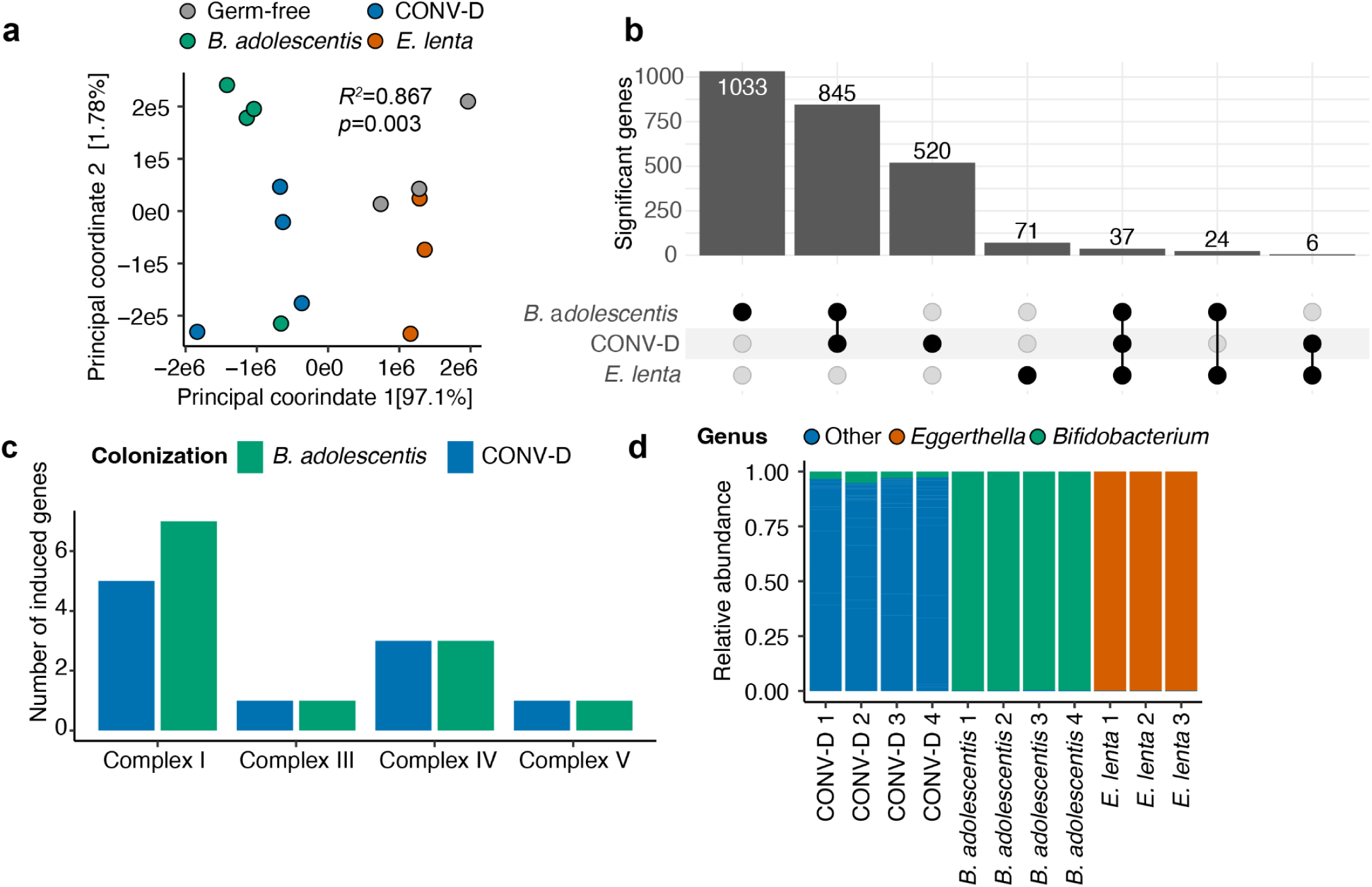
The lung transcriptome is responsive to gut microbiome colonization. **(a)** Principal coordinate analysis of bulk RNA sequencing from lung tissue isolated from GF, *B. adolescentis* colonized mice, conventionalized (CONV-D) mice, or *E. lenta* colonized mice. Principal coordinate analysis with PERMANOVA testing *R^2^* and *p*-value annotated between all four colonization states. **(b)** Differentially expressed genes between *B. adolescentis* colonized mice, CONV-D mice, and *E. lenta* colonized mice and the GF group. **(c)** Differentially stimulated electron transport chain genes by transport complex. **(d)** Relative abundance of 16S-Seq of indicated mice colored by genus of *Bifidobacterium*, *Eggerthella*, or all other genera (Other). n=3 GF, n=4 CONV-D, n=4 *B. adolescentis* mice, and n=3 *E. lenta* mice for all panels.

**Extended Data Figure 5:**
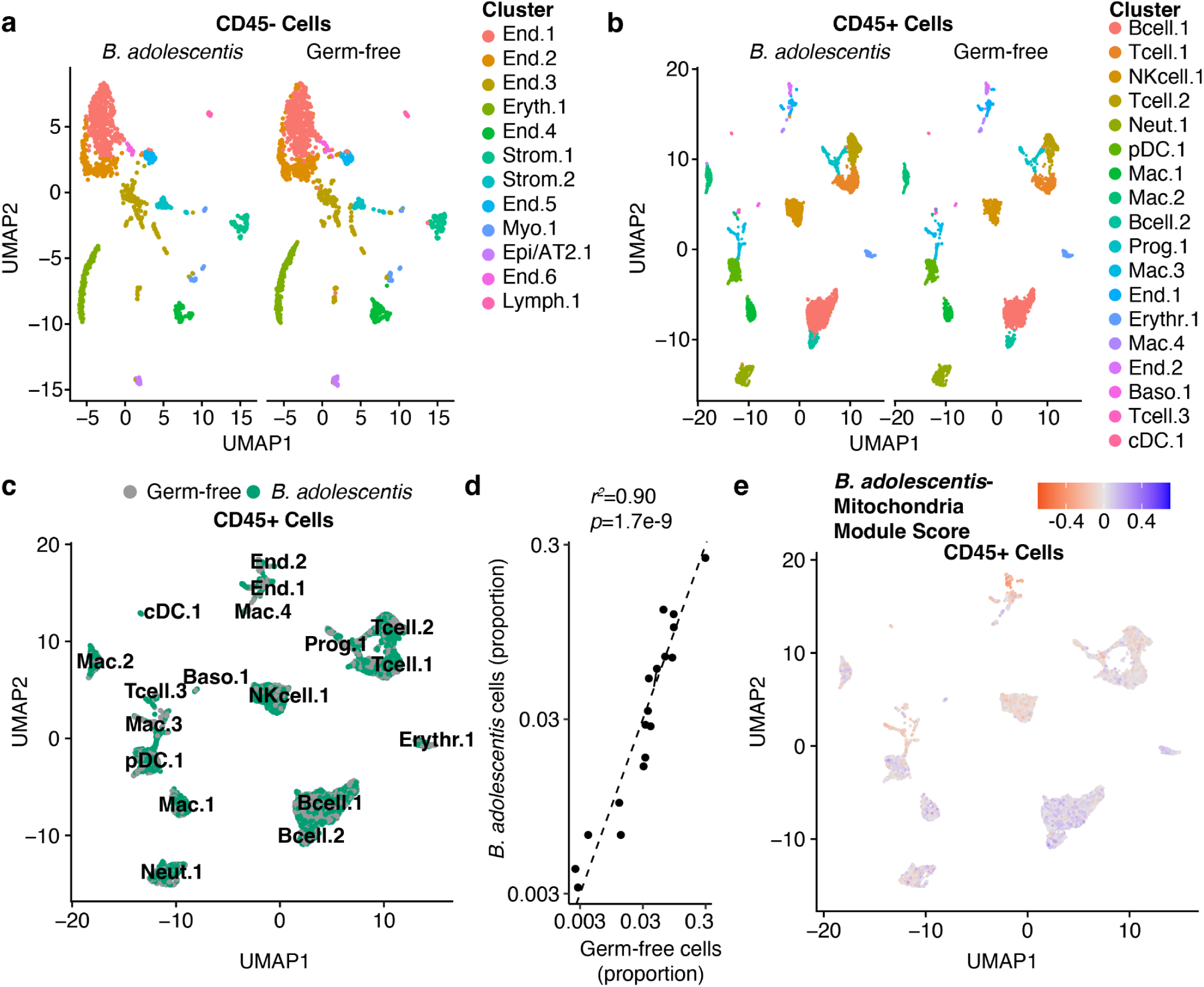
*B. adolescentis* does not significantly impact the mitochondrial activity of CD45+ cells in the mouse lung. **(a-b)** Cluster assignment of scRNA-seq of live CD45-**(a)** or CD45+ **(b)** cells from a representative *B. adolescentis* colonized or GF mouse. Clusters assignments by Seurat were annotated with the Mouse Cell Atlas. **(c)** Individual cells are colored by *B. adolescentis* colonized or GF groups. **(d)** The proportion of each cluster in the *B. adolescentis* colonized vs GF mice is plotted. A Pearson’s *r^2^* and *p*-value are annotated above the plot. **(e)** A module score was calculated in Seurat from mitochondrial genes significantly impacted by *B. adolescentis* colonization (adjusted *p*<0.1), and cells are colored by module score.

**Extended Data Figure 6:**
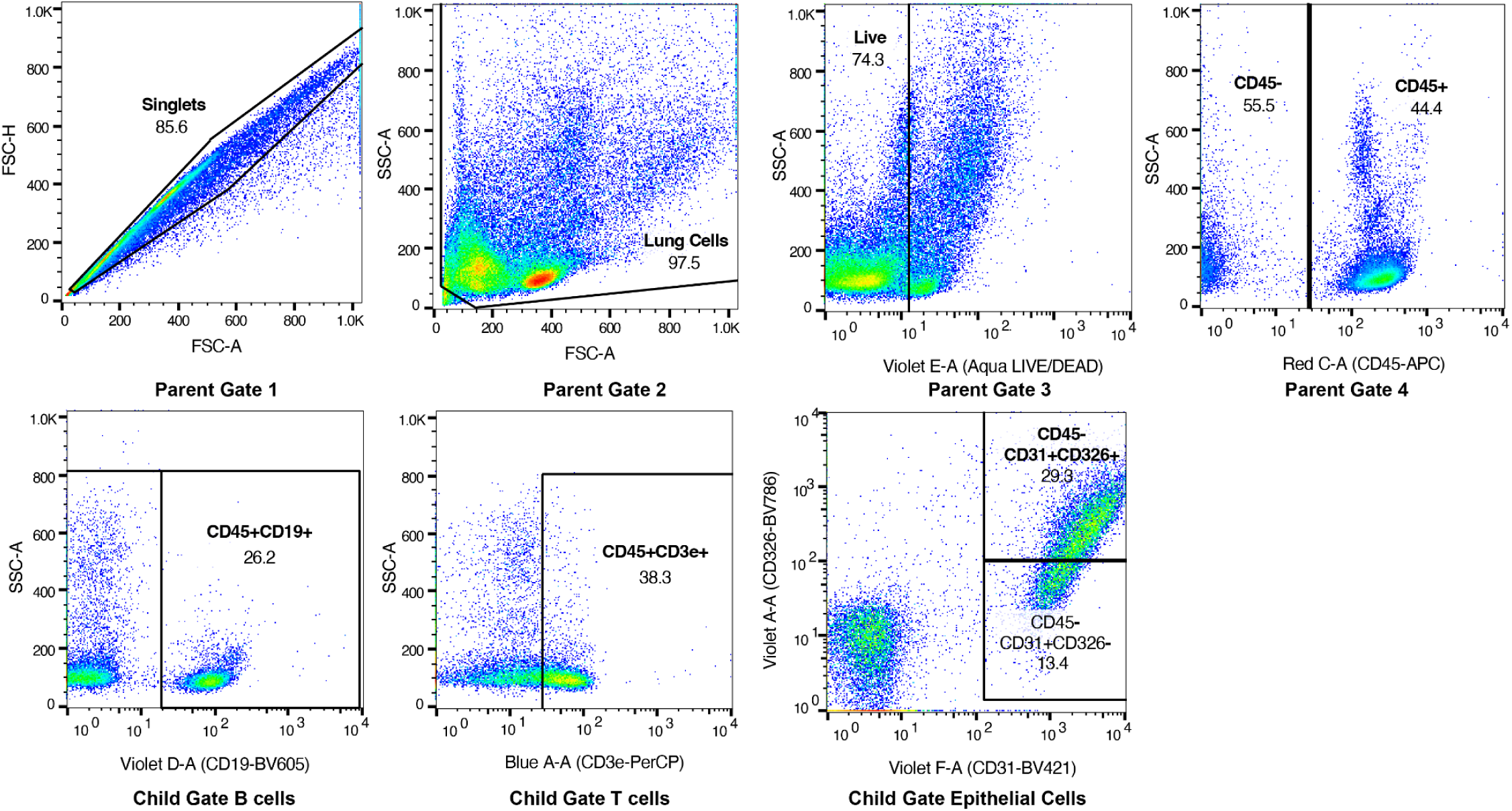
Gating strategy for pulmonary epithelial cells and flow cytometry. Parent gates were set on Singlets (Parent Gate 1), followed by Lung Cells (Parent Gate 2), followed by Live Cells (Parent Gate 3), followed by CD45+ or CD45-cell types (Parent Gate 4). The main text shows Child Gate Epithelial Cells which is from Parent Gate 4.

**Extended Data Figure 7:**
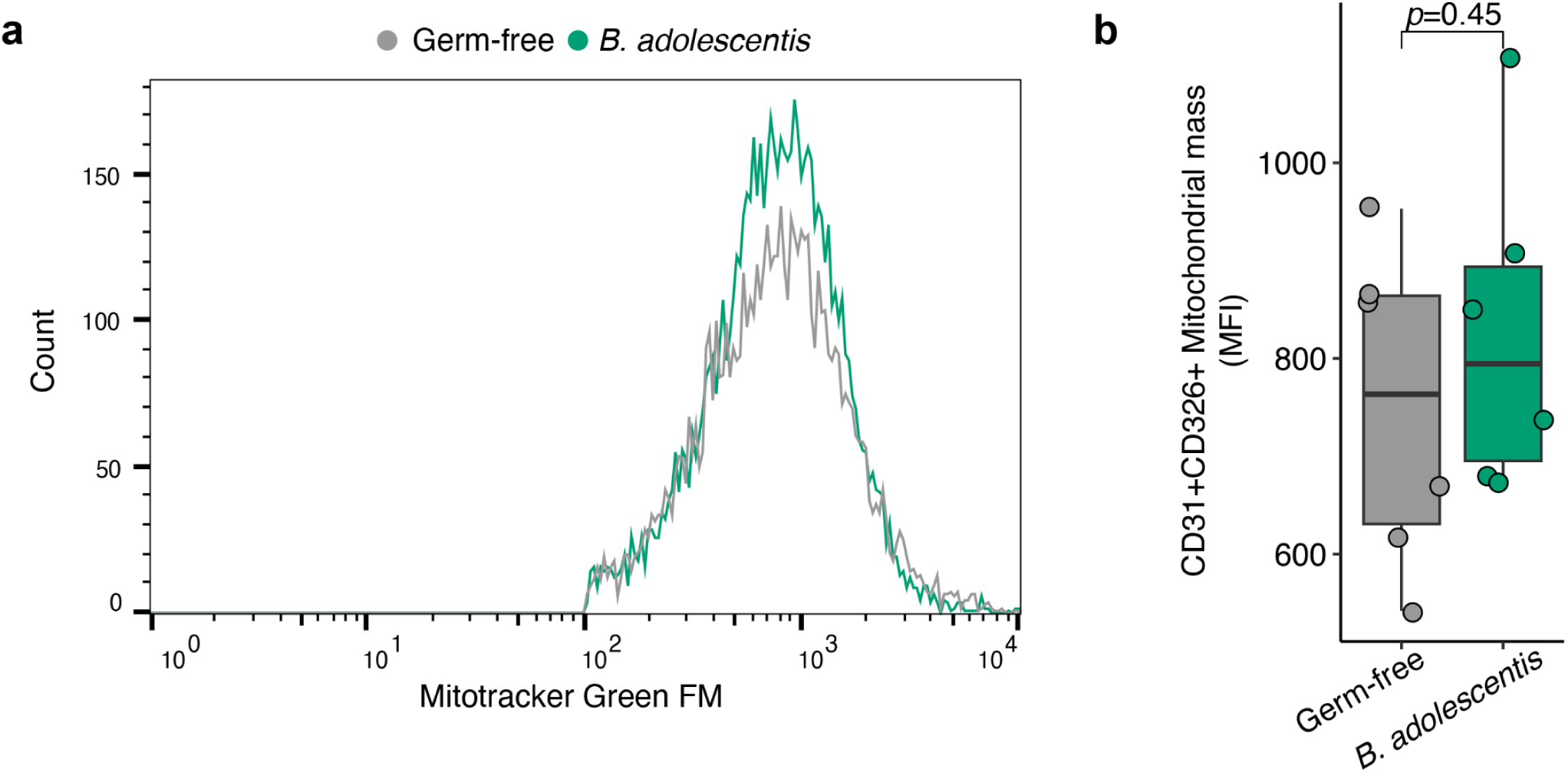
*B. adolescentis* does not increase mitochondrial biomass of lung epithelial cells. **(a)** Histogram of the MitoTracker Green probe gated on epithelial child gate from **Extended Fig. 6**. **(b)** Mean fluorescence intensity (MFI) of MitoTracker Green probe which indicates mitochondrial biomass. n=6 mice per group.

**Extended Data Figure 8:**
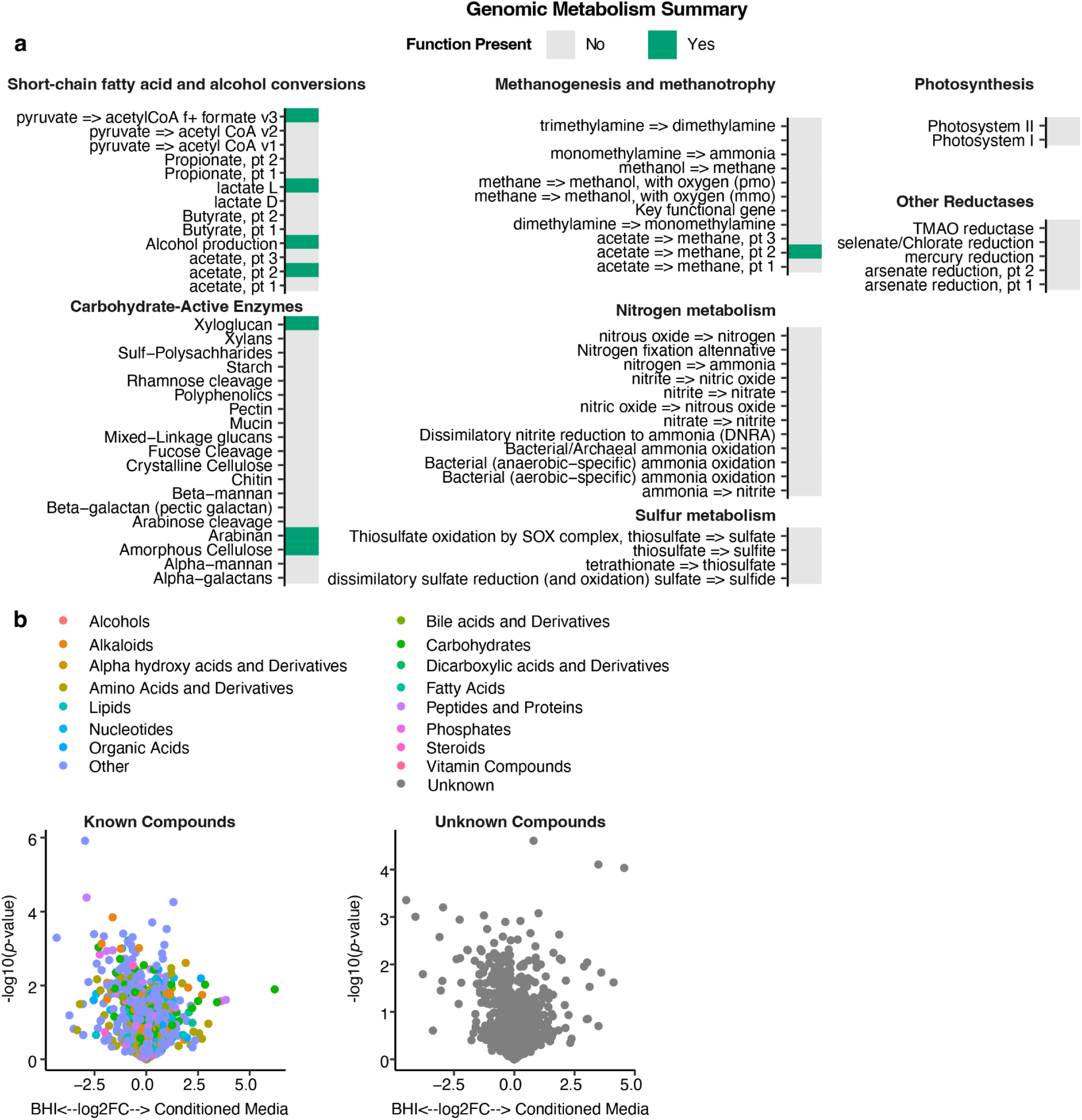
The genomic and metabolic capacity of *B. adolescentis*. **(a)** The full-length *B. adolescentis* genome was input to the Distilled and Refined Annotation of Metabolism (DRAM v0.1.2) tool. The outputted metabolism summary indicates whether and which specific metabolic conversions are predicted to be present within the genome of *B. adolescentis*. **(b)** Untargeted metabolomic profiling of *B. adolescentis* conditioned media compared to BHI. Compound category is indicated by point color. Compounds are separated based on whether compounds could be annotated by MSDIAL (Known) or could not be annotated (Unknown). n=3 independent conditioned media specimens per group.

**Extended Data Figure 9:**
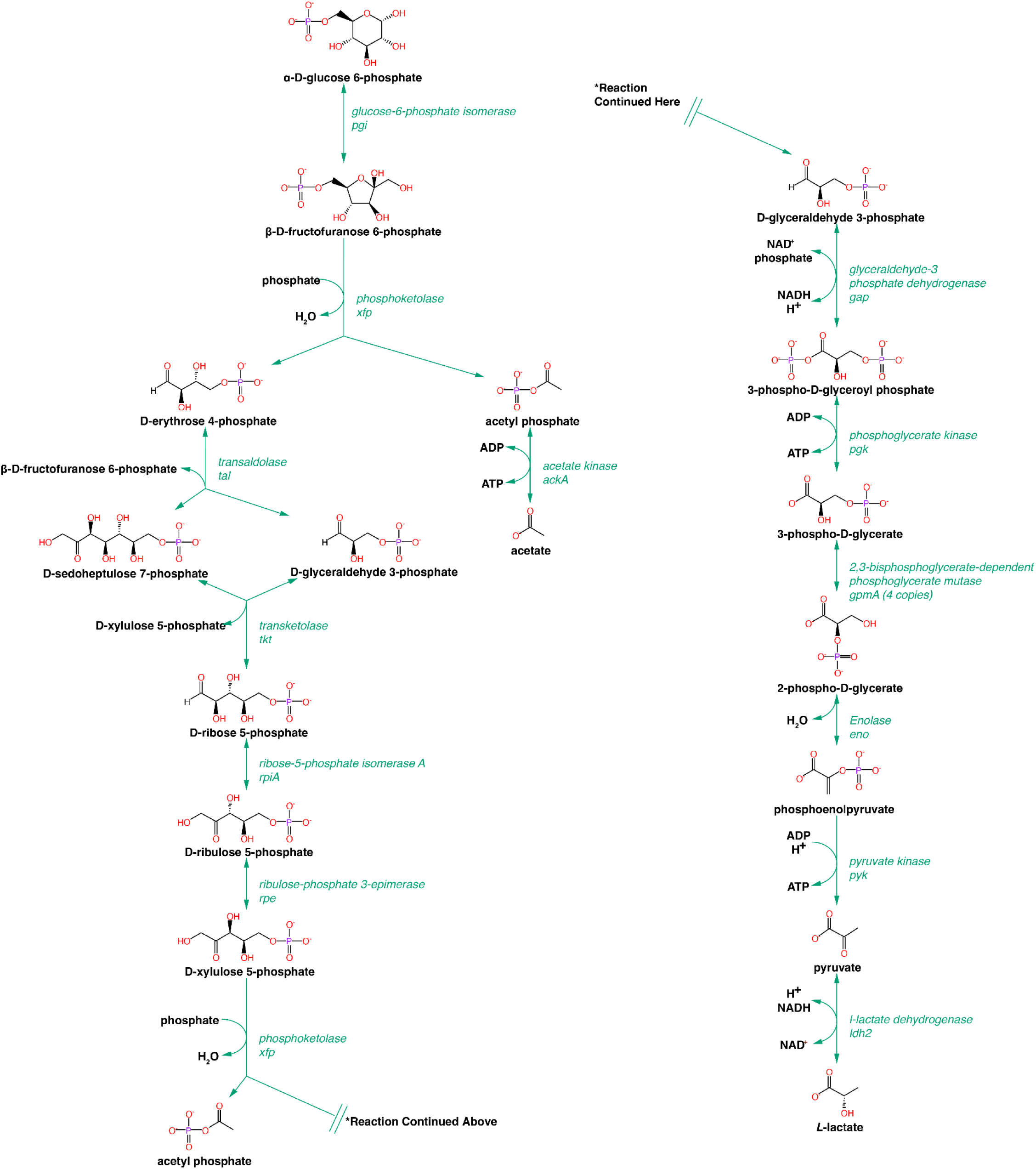
Complete *L*-lactate production pathway in *B. adolescentis* BD1. The *L-*lactate production pathway in *B. adolescentis* BD1 was constructed using MetaCyC. All genes in green are genes annotated within the *B. adolescentis* BD1 genome and can be found in Supplementary Table 3b. Starting, intermediate, and end products are shown. Other bifidobacteria contain a 2,3-bisphosphoglycerate-independent phosphoglycerate mutase (gpmM), and *B. adolescentis* BD1 maintains four copies of the 2,3-bisphosphoglycerate-independent phosphoglycerate mutase (gpmA), which catalyzes the same reaction with lower enzymatic specificity. The double break in the diagram indicates where the reaction continues to facilitate viewing on a standard page.

**Extended Data Figure 10:**
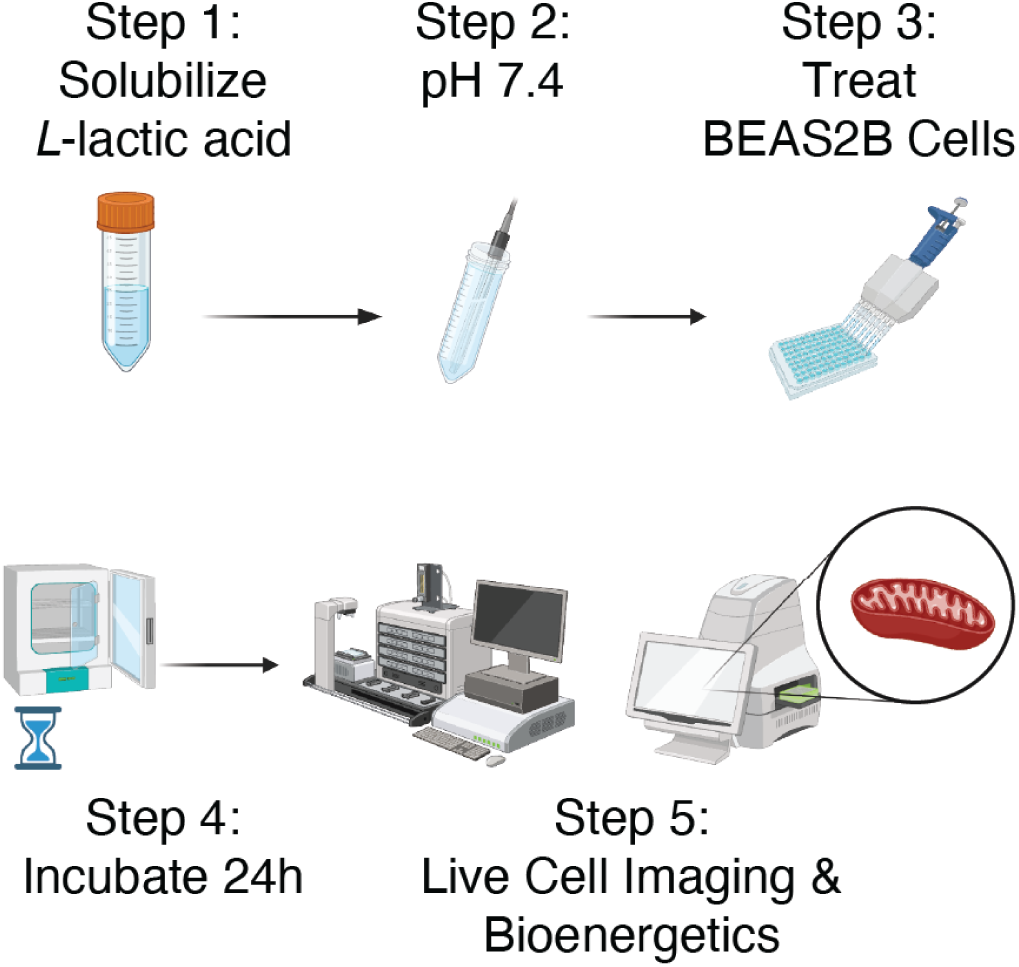
Workflow to profile the impact of metabolites on lung epithelial activity. Workflow for testing the impact of gut bacterial metabolites on lung epithelial cells *in vitro*. *L*-lactate was solubilized in water. It was then pH adjusted to 7.4. BEAS-2B cells were treated with the indicated doses of *L-*lactate for 24 hours, with subsequent live cell imaging and bioenergetic assessment. Created with Biorender.com.

